# Secretory carrier membrane proteins regulate aquaporin trafficking in Arabidopsis

**DOI:** 10.1101/2025.07.03.662988

**Authors:** Qihang Jiang, Omar Hdedeh, Michaël Vandorpe, Ana Romina Fox, Hongyan Liu, Lei Ding, Mattias Vermeersch, Evelien Mylle, Jonah Nolf, Alvaro Furones Cuadrado, Julia Kraus, Dominique Eeckhout, Daniela Kocourková, Tereza Korec Podmanická, Thomas B. Jacobs, Jonathan Michael Dragwidge, Bert De Rybel, Ive De Smet, Roman Pleskot, François Chaumont, Daniël Van Damme

## Abstract

Aquaporins belonging to the plasma membrane intrinsic protein (PIP) subfamily are channel proteins that control water flow across cells, allowing plants to rapidly adjust hydraulic conductivity and thereby sustain growth, gas exchange, and recovery from drought. We show that secretory carrier membrane proteins (SCAMP) regulate the abundance of the aquaporins and thereby water transport in Arabidopsis root cells. SCAMPs are evolutionarily conserved multi-spanning transmembrane proteins. In animal cells, they function in secretion, endocytosis and autophagy. Knowledge on their role in plants is restricted to localization and trafficking experiments in heterologous systems and few genetic perturbation experiments. Here, we analysed all five members of the Arabidopsis SCAMP family. We identified conserved tyrosine motifs assisting in transport to the plasma membrane and N-terminally located NPF motifs that are required for internalization. SCAMPs dimerize both at the plasma membrane and endosomes, and dimerization is required for their internalization. Functionally, several PIPs were identified as common targets of multiple SCAMP isoforms. Triple and quintuple *scamp* mutants show mild developmental delay under standard growth conditions, but they are less sensitive to drought-induced leaf wilting. This observation cannot be explained by altered stomatal dynamics or densities, nor by differential soil water content. However, *scamp* mutant root protoplasts contain less PIPs and swell less under hypotonic conditions compared to wild type, indicating reduced PIP plasma membrane levels. In conclusion, our research identifies the SCAMP membrane trafficking proteins as regulators of PIP abundance at the plasma membrane in root cells. We propose that the reduced PIP levels in the *scamp* mutants act as a priming mechanism, allowing them to respond better to conditions of reduced water availability.

## Introduction

Secretory carrier membrane proteins (SCAMPs) constitute a family of integral membrane proteins that are highly enriched in the secretory membranes of both animal and plant cells (Hubbard et al. 2000; Law et al. 2012). SCAMPs contain four transmembrane domains (TMDs) that are evolutionarily conserved (Law et al. 2012), as well as short loops between them and cytosolic tails at both N- and C-termini that contain linear trafficking motifs. Based on their tissue distribution and localization to secretory membranes, SCAMPs were proposed to be involved in vesicle-mediated transport(Brand et al. 1991). Animal cells have five SCAMPs, with SCAMP1-4 exhibiting a widespread expression and SCAMP5 being primarily expressed in neuronal tissues(Han et al. 2009; Zhao et al. 2014). Several studies in animal cells have underscored SCAMPs intracellular localization and regulation of post-Golgi trafficking. SCAMPs have been implicated in the trafficking of the (Na^+^, K^+^)/H^+^ exchanger (NHE7) between the trans-Golgi network (TGN) and recycling vesicles (Lin et al. 2005). They can exert both positive and negative effects on cargo exocytosis to the plasma membrane (PM) (Diering et al. 2009; Fjorback et al. 2011; Zaarour et al. 2011) and participate in forming fusion pores on the PM by interacting with phosphatidylinositol 4,5-bisphosphate(Liao et al. 2007; Fjorback et al. 2011). Additionally, SCAMPs regulate the frequency of vesicle fusion events (Liao et al. 2008) and contribute to fusion during exocytosis (Guo et al. 2002; Liu et al. 2002). SCAMPs also play a role in endocytosis (Fernández-Chacón and Südhof TC 2000; Zhao et al. 2014), protein recycling (Aoh et al. 2009), autophagy (Yang et al. 2017; Hama et al. 2023) and vesicle protein clearance from the synaptic release site (Park et al. 2018). Consistent with their roles in secretory vesicle trafficking and cell-cell adhesion, SCAMP deficiencies have been implicated in various human cancers (Zhang et al. 2017; Asghari et al. 2018; Vadakekolathu et al. 2018), as well as neurological disorders (Hubert et al. 2020; Jiao et al. 2020).

In plants, our understanding of SCAMP function is very limited. Next to their topology, plant and animal SCAMPs share some trafficking related motifs, although many plant SCAMPs contain an additional tyrosine-based sorting motif, which is not present in their animal counterparts (Law et al. 2012). Furthermore, SCAMPs from animals and plants cluster in separate branches, which suggests that SCAMPs evolved independently in these kingdoms (Law et al. 2012). Localization studies in tobacco bright yellow-2 (BY-2) cells revealed the presence of rice SCAMP1 at both the PM and endosomes, and its trafficking was found to be sensitive to Brefeldin A and Wortmannin (Sheung et al. 2008; Cai et al. 2011). Tobacco SCAMP2 is also localized to the TGN/early endosome (EE), the cell plate, and the PM. Wortmannin and Brefeldin A treatment did not cause a similar relocalization of this SCAMP compared to rice SCAMP1 (Toyooka et al. 2009), implying that individual plant SCAMP proteins may have specific localisation patterns and functions.

As one of the five SCAMP family members in Arabidopsis, SCAMP5 was found as an interactor of the endocytic TPLATE complex (TPC) via proximity biotinylation (Arora et al. 2020). More specifically, the double NPF repeat in the N-terminus of Arabidopsis SCAMP5 interacts with the first Eps15 homology (EH) domain of the AtEH1/Pan1 TPC subunit (Yperman et al. 2021). A truncated construct of SCAMP5, lacking the N-terminal domain accumulates at the PM (Yperman et al. 2021). This data suggests that SCAMPs contain distinct motifs that control their trafficking, although the distinct anterograde and retrograde trafficking pathways that are affected are unknown.

In Arabidopsis, *scamp2, scamp3, scamp4* and *scamp5* single mutants were reported to exhibit different degrees of insensitivity to abscisic acid (ABA), flg22, chitin, CaCl_2_ or H_2_O_2_-mediated stomatal closure, suggesting a role for SCAMPs in mediating stress responses (Bourdais et al. 2019). Few papers report on specific deficiencies associated with altered *SCAMP* levels in other plants. Silencing *SCAMP* abolishes pollen tube growth in lily (*Lilium longiflorum*) (Wang et al. 2010) and RNAi-mediated downregulation of *SCAMP3* and *SCAMP6* in poplar increases the secretion of secondary cell wall components in the stem (Obudulu et al. 2018). In soybean, cotton and rubber tree, expression of *SCAMP* family members was reported to change in response to abiotic stress conditions. Downregulation of *SCAMP2* and S*CAMP4* in cotton rendered plants more sensitive to salt stress (He et al. 2024), while over expression of *SCAMP5* in soybean rendered plants more tolerant to salt stress and this salt tolerance phenotype was correlated with reduced sodium accumulation in the plants (Wang et al. 2023). On the other hand, overexpression of rubber tree *SCAMP3* in poplar was shown to significantly promote plant height (Yang et al. 2024). The above observations link SCAMPs to various aspects of plant physiology, but because our current knowledge on the role of this protein family is very fragmented and indirect, we cannot pinpoint the subcellular functions that are causal to the observations made.

Here, we characterized the SCAMP family in Arabidopsis. We identified motifs involved in anterograde and in retrograde trafficking. Higher-order mutants show minor defects in root and rosette development. Notably, we discovered a direct connection between SCAMPs and the aquaporins belonging to the plasma membrane intrinsic protein (PIP) subfamily. We found that both PIP1 and PIP2 abundance was lower in triple and quintuple *scamp* mutant roots compared to wild type. In contrast, PIP2 levels were increased in *scamp* mutant leaves, while PIP1 levels remained similar to those in control cels. We show also that *scamp* mutant root protoplasts display reduced water transport capacity. Our results indicate a complex role for SCAMPs in differentially modulating PIP trafficking or stability in Arabidopsis above- and belowground organs and we hypothesize that the altered PIP levels contribute to the observed drought tolerance phenotype of the *scamp135* mutants.

## Results

### SCAMP5 localizes at the PM and the TGN/EE in Arabidopsis root meristems

The *SCAMP* gene family in Arabidopsis comprises five members: *SCAMP1 (At1g61250)*, *SCAMP2 (At1g11180)*, *SCAMP3 (At2g20840)*, *SCAMP4 (At1g03550)* and *SCAMP5 (At1g32050)*. We initially expressed all *SCAMPs* in Arabidopsis as genomic fusion proteins C-terminally tagged with enhanced green fluorescent protein (eGFP), under the control of their respective native *SCAMP* promoters (Supplementary Figure 1A). Analysis of our transgenic lines, alongside published single-cell RNA sequencing data (Wendrich et al. 2020; Berrío et al. 2022; Kim et al. 2023) (Supplementary Figure 1B), demonstrated that SCAMP1, 3, and 5 are widely expressed in various tissues and that they localize at the PM, endosomes, and forming cell plates. SCAMP2 is weakly expressed, and the SCAMP2 localization pattern in the root meristem and the stomata is reminiscent of that of the ER. scRNAseq data revealed *SCAMP4* expression to be very low to absent in root and leaf tissues, alongside some expression that is restricted to mature guard cells. The very low expression of *SCAMP4* was also evident from our confocal imaging (Supplementary Figure 1).

Given the widespread and high expression of *SCAMP5,* we initially focused on this family member. SCAMP5 localises to the PM, cell plate, and endosomes in epidermal root tip cells (Figure 1A). Colocalization and Brefeldin A (BFA) experiments revealed that its endosomal localization predominantly overlaps with the TGN/EE as comparing Golgi, TGN and MVB markers, the highest degree of endosomal colocalization was observed with the VHA-a1 TGN marker. Moreover, both SCAMP5 and VHA-a1 also perfectly overlapped at BFA bodies (Figure 1B and 1C).

**Figure 1.**
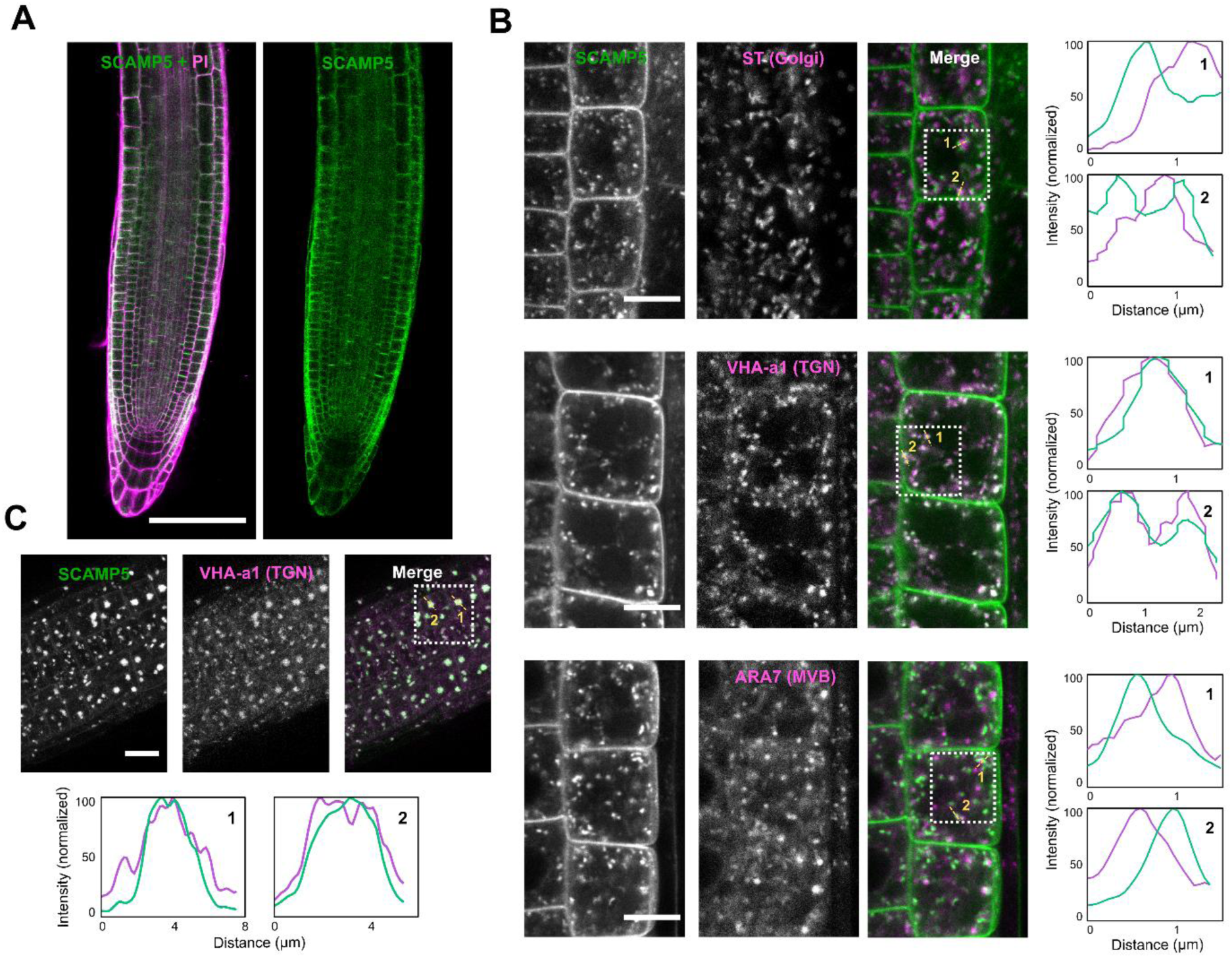
SCAMP5 localizes at the PM and the TGN/EE in Arabidopsis root meristems. **A)** Representative confocal image of a propidium iodide (PI) stained root tip expressing SCAMP5-GFP. Scale bar, 100 µm. **B)** Colocalization analysis of SCAMP5-GFP with VHA-a1-RFP (TGN/EE marker), ST-RFP (Golgi marker), and RFP-ARA7 (late endosome marker). Scale bar, 10 µm. **C)** Colocalization of SCAMP5-GFP with VHA-a1-RFP after brefeldin A (BFA) and cycloheximide (CHX) treatment. The seedlings were pre-treated in liquid 1/2 MS medium with 50 µM CHX for 30 min and then treated in liquid 1/2 MS medium with 50 µM BFA and CHX for 60 min. Scale bar, 20 µm. This experiment was repeated twice. The graphs in panel B and C represent multi-plot visualizations of the colocalization of the indicated positions in the dotted squares.

### The double NPF motif controls SCAMP endocytosis

The localization pattern of SCAMP5 at the PM and TGN/EE suggests that it traffics between these compartments. Trafficking of membrane proteins within the endomembrane system can be regulated by sorting motifs, which are recognized by adaptor protein (AP) complexes (Arora and van Damme 2021). Truncation experiments previously revealed that the double NPF motif (N=asparagine, P=proline, F=phenylalanine) regulates SCAMP5 endocytosis through a direct interaction with the TPC subunit AtEH1/Pan1 (Yperman et al. 2021). The N-termini of SCAMP1, 4, and 5 contain the double NPF motif, while SCAMP2 and 3 only have a single NPF motif (Supplementary Figure 1C). Additionally, multiple conserved tyrosine motifs (YXXϕ; Y = tyrosine, ϕ = a bulky hydrophobic residue), known to interact with APs, were identified between transmembrane domains 2 (TMD2) and 3, as well as in the C-terminal tail of all SCAMPs. In contrast, an acidic dileucine motif ([D/E]xxxL[L/I]; D = aspartic acid, E = glutamic acid, L = leucine, I = isoleucine), facilitating interaction with the small subunit of the AP complexes in animal cells (Kelly et al. 2008), was found exclusively in SCAMP4 (Supplementary Figure 1C). Given the poor conservation of the acidic dileucine motif, we focused on the roles of the NPF and tyrosine motifs present in the cytoplasmic regions of all SCAMPs (Figure 2A).

**Figure 2.**
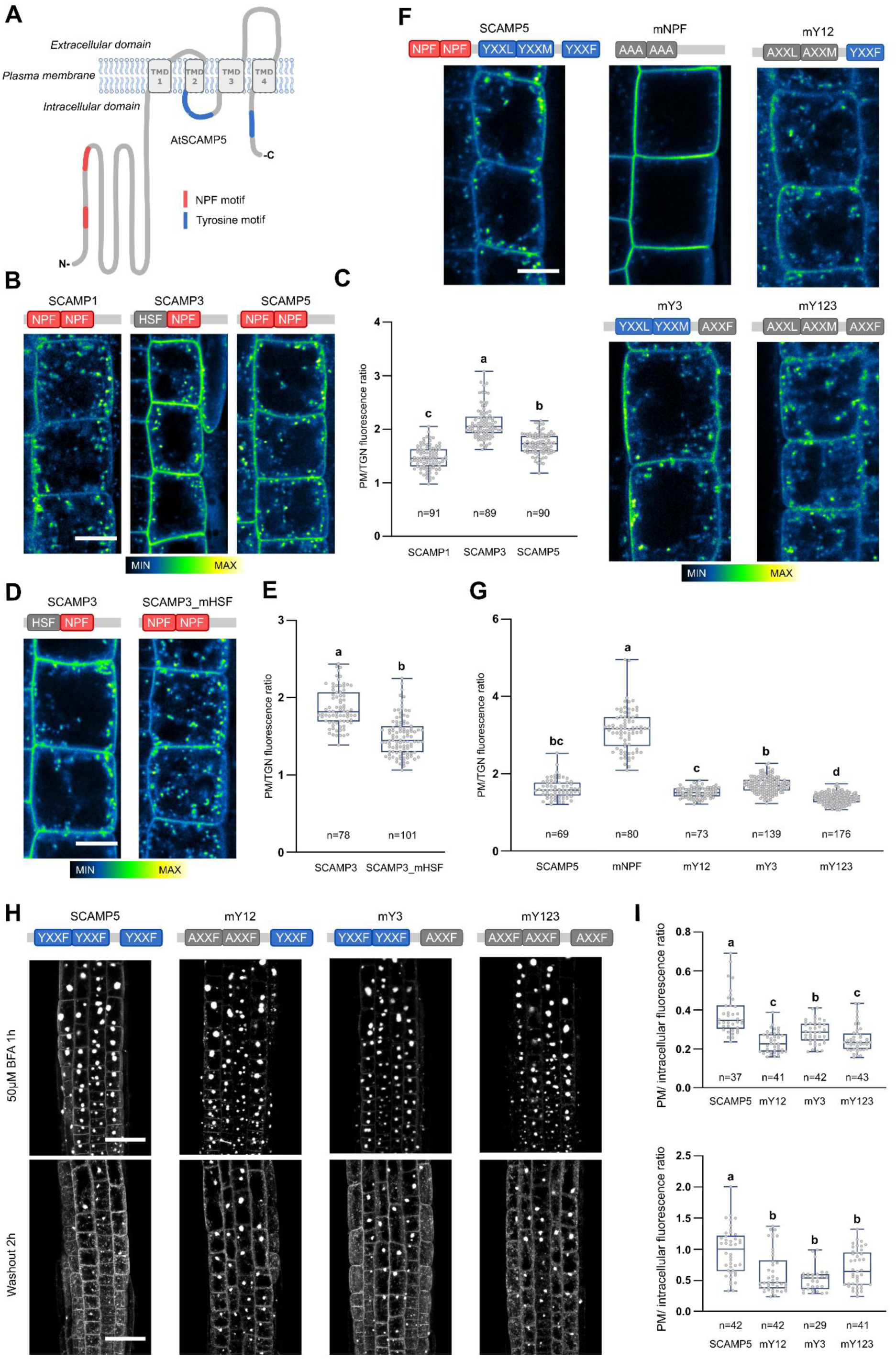
The tyrosine and the double NPF motifs control anterograde transport and endocytosis of SCAMP proteins. **A)** Schematic representation of the secondary protein structure of SCAMP5 as predicted by Protter (Omasits et al. 2014). The NPF and the tyrosine motifs are represented by red and blue boxes respectively. TMD: transmembrane domain. N, C: N- and C-terminus of SCAMP5. **B, D and F)** Representative intensity coloured confocal images of root epidermal cells expressing SCAMP1, SCAMP3, and SCAMP5 GFP fusions as well as motif-mutated forms of SCAMP3 and SCAMP5. SCAMP fusions in panel B and D are in the Col-0 background. SCAMP5 fusions in panel F are in the *scamp5-1* mutant background Scale bar, 10 µm. The presence or absence of the respective double NPF motifs (red/grey) or the tyrosine motifs (blue) are shown above the images. **C, E and G)** Box plot graph of the PM over TGN/EE intensity ratios of the constructs from panels B, D and F. The numbers at the bottom represent the number of cells analysed. Quantifications were done on at least 12 seedlings (C), at least 7 seedlings (D) or at least 10 T2 generation seedlings from two independent lines (G) and the data of both lines was merged. Different letters in panel C and G represent significant differences evaluated by a one-way ANOVA test (P<0.0001). The experiments in C and G were repeated twice. Different letters in panel E represent significant differences of a Student’s t-test (P<0.0001). This experiment was performed once. For each box plot, the bottom and top edges indicate the 25th and 75th percentiles. The central line is the median and whiskers extend to the full data range. **H)** Representative confocal images of SCAMP5 and motif-mutated SCAMP5 isoforms following BFA treatment (top row) and following BFA washout (bottom row). The presence or absence of the respective tyrosine motifs (blue/grey) is shown above the images. Scale bar, 30 µm. I) Box plot graph of the PM over intracellular intensity ratio of SCAMP5 and the motif-mutated SCAMP5 isoforms following BFA treatment (top) and following BFA washout (bottom). The numbers at the bottom represent the number of cells analysed from at least 7 seedlings. Different letters represent significant differences evaluated by a one-way ANOVA test (P<0.05). For each box in the box plot, the bottom and top edges indicate the 25th and 75th percentiles. The central line is the median and whiskers extend to the full data range. This experiment was performed once. mHSF: mutated HSF to NPF; mNPF: both NPF motifs mutated to AAA; mY12: first and second tyrosine motifs mutated to AXXF; mY3: third tyrosine motif mutated to AxxF; mY123: all tyrosine motifs mutated to AXXF.

SCAMP1, 3, and 5 show abundant expression in root cells but differ in the amount of NPF motifs in their N-terminus (Supplementary Figure 1C). SCAMP3 exhibited a stronger localization at the PM compared to SCAMP1 and SCAMP5, suggesting a lower rate of internalization (Figure 2B and 2C). Given that SCAMP3 only possesses a single NPF motif while SCAMP1 and SCAMP5 each have two, we hypothesized that this might be causal to the observed localization pattern. To test this hypothesis, we mutated H7S8F9 in SCAMP3 to N7P8F9 to generate an artificial double NPF motif. The mutated protein (SCAMP3_mHSF) was tagged with GFP, under the control of its native promoter, and expressed in the Col-0 background. Compared to the genomic SCAMP3, SCAMP3_mHSF exhibited an increased endosomal localization, indicating enhanced internalization (Figure 2D and 2E). The importance of the double NPF motif for SCAMP internalization was further confirmed by constructing SCAMP5_mNPF (N8A; P9A; F10A, N18A; P19A; F20A). SCAMP5_mNPF, expressed in the *scamp5-1* mutant background (Supplementary Figure 2), was almost exclusively located at the PM, and the PM/TGN ratio was significantly higher compared to the genomic SCAMP5 in the same genetic background (Figure 2F and 2G). In conclusion, the relocalization of SCAMP3_mHSF as well as the blocked internalization of SCAMP5-mNPF implies that the double NPF motif of SCAMP5 is the predominant endocytic motif.

### The tyrosine motifs aid anterograde trafficking of SCAMP5 from the TGN/EE

The role of the three tyrosine motifs in SCAMP trafficking was assessed by generating mutated SCAMP5 isoforms targeting these motifs (Figure 2A and 2F). To explore potential differences in the functions of the tyrosine motifs, we mutated them in different combinations: SCAMP5_mY12(Y_170_A, Y_174_A), SCAMP5_mY3(Y_257_A), and SCAMP5_mY123(Y_170_A, Y_174_A, Y_257_A). SCAMP5_mY123, lacking all tyrosine motifs, exhibited a decreased ratio of PM over TGN/EE fluorescence, while disruption of one or two motifs did not significantly affect SCAMP5 localization (Figure 2F and 2G). To verify the role of the tyrosine motifs in post-TGN trafficking, we subjected the tyrosine motif-mutated SCAMP5 isoforms to a BFA washout experiment. Removing BFA resumes exocytosis and therefore allows assessing the recycling potential of endosomal proteins. All tyrosine motif mutated isoforms had a decreased ratio of PM over BFA body intensity following BFA treatment, and showed a longer retention time at the TGN/EE following washout compared to the control (Figure 2H and 2I), suggesting that these tyrosine motifs function in post-Golgi trafficking of SCAMP5.

Plants lacking SCAMP5 are phenotypically similar to wild type at the young seedling stage (Supplementary Figure 2). Still, functional testing revealed that *scamp5-1* mutants expressing SCAMP5_mNPF isoforms showed reduced primary root and dark-grown hypocotyl length compared to the *scamp5-1* mutants or *scamp5-1* mutants expressing SCAMP5_mY123 (Supplementary Figure 2). These mild dominant negative effects suggest a role of SCAMP5 in facilitating endocytic cargo trafficking and/or dimerization-dependent internalization.

### Arabidopsis SCAMPs function as dimers

SCAMPs are known to dimerize in mammalian systems (Wu and David Castle 1998; Lin et al. 2005). We therefore tested whether this feature is conserved for the Arabidopsis SCAMPs. Ratiometric bimolecular fluorescence complementation (rBiFC) revealed homo-dimerization of SCAMP5 and hetero-dimerization of SCAMP5 and SCAMP3 (Figure 3A-B). As a control, we included the hetero-dimerization between SCAMP5 and another PM localized transmembrane protein, the brassinosteroid receptor BRI1. To expand our binary interaction analyses, we predicted all possible SCAMP dimers using AlphaFold2 (Supplementary Figure 3A). All combinations generated models with high confidence scores, and the predicted aligned error (PAE) graphs revealed a dominant role for the N-terminal helix in SCAMP dimerization. Yeast two-hybrid (Y2H) analysis between the N-terminal regions of all SCAMPs confirmed that homo- and heterodimerization of SCAMPs occurs via the N-terminal alpha helical domain (Supplementary Figure 3B).

**Figure 3.**
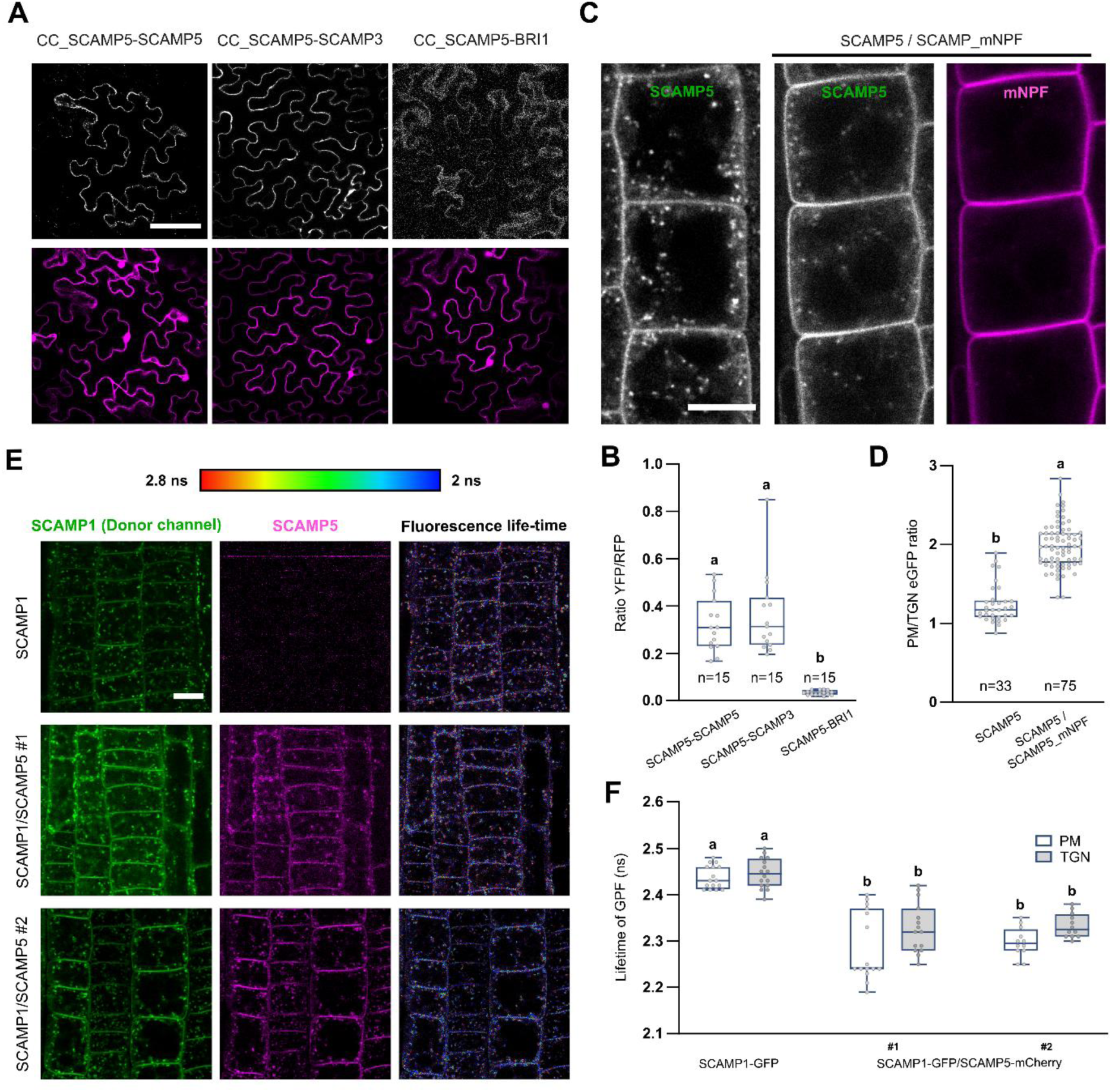
SCAMPs homo- and hetero-dimerize. **A and B)** Ratiometric BiFC analysis and box plot quantification (B) showing homo-dimerization of SCAMP5 dimerization between SCAMP5 and SCAMP3. CC_SCAMP5-BRI1 was used as a negative control. CC refers to the C-terminal fusion of the nYFP and cYFP halves to the SCAMPs. Scale bar, 50 µm. The numbers at the bottom represent the number of cells analysed. Different letters represent significant differences evaluated by a one-way ANOVA test (P<0.0001). This experiment was performed once. **C and D)** Representative confocal images of Arabidopsis Col-0 root cells expressing SCAMP5-GFP or co-expressing SCAMP5-GFP and SCAMP5_mNPF-mCherry and box plot graph of the PM over TGN/EE intensity of SCAMP5-GFP from the lines in panel B (D). Scale bar, 10 µm. The numbers at the bottom represent the number of cells analysed from at least 7 seedlings. Different letters represent significant differences evaluated by the Student’s t-test (P<0.0001). This experiment was repeated twice. **E and F)** FRET-FLIM analysis of SCAMP1 and SCAMP5 interaction in planta. From top to bottom, representative images of root tips, expressing SCAMP1-GFP and two independent lines co-expressing SCAMP1-GFP and SCAMP5-mCherry. The donor (GFP) fluorescence life-time calibration corresponds to the adjacent scale bar. The bar chart in F shows the donor fluorescence life-time quantifications of the SCAMP1-GFP line and the two independent Arabidopsis lines (#1 and #2) co-expressing SCAMP5-mCherry and SCAMP1-GFP. The PM region (white box) and the TGN/EE region (grey box) were quantified separately. Each dot on the graph represents the quantification of an individual seedling, which represents the measurement of +/- 3 cells. More than 10 seedlings were quantified for each line. Different letters represent significant differences evaluated by a one-way ANOVA test (P<0.001). This experiment was repeated twice. For each box plot, the bottom and top edges indicate the 25th and 75th percentiles. The central line is the median and whiskers extend to the full data range.

To test whether dimerization played a role in the trafficking of the SCAMP proteins, we transformed SCAMP5-GFP expressing Arabidopsis lines with SCAMP5_mNPF-mCherry (Figure 3C). SCAMP5-GFP signal was more prominent at the PM when co-expressed with SCAMP5_mNPF-mCherry, suggesting that homodimerization precedes SCAMP5 endocytosis (Figure 3D). To test if heterodimerization also occurred and whether this was specific for the PM or not, we performed Förster resonance energy transfer fluorescence life-time imaging microscopy (FRET-FLIM) experiments using Arabidopsis seedlings expressing SCAMP1-GFP, or co-expressing SCAMP5-mCherry and SCAMP1-GFP (Figure 3E). This analysis revealed a reduced fluorescence life-time of the SCAMP1-GFP donor in combination with the SCAMP5-mCherry acceptor both at the PM and at the TGN/EE (Figure 3F), showing that SCAMP5 heterodimerization occurs at both locations.

### SCAMP1, 3 and 5 interact with several members of the aquaporin PIP subfamily

Mammalian SCAMPs facilitate cargo trafficking (Wu and David Castle 1998; Lin et al. 2005; Lee et al. 2021). To identify putative cargo proteins for Arabidopsis SCAMPs, we conducted GFP pull-down assays followed by mass spectrometry analyses using Arabidopsis seedlings expressing GFP fusions of SCAMP1, SCAMP3 and SCAMP5 expressed under their endogenous promotors (Supplementary Figure 4). In addition to confirming SCAMP heterodimerization, many type 1 and type 2 PIP aquaporin members were significantly enriched and common to all three SCAMP isoforms (Figure 4A and Supplementary Table 2). PIPs serve as primary water channels in the PM, mediating water balance with the environment (Chaumont and Tyerman 2014). The interaction between SCAMP5 and four PIP2s and one PIP1 was confirmed via the split ubiquitin system (SUS) in yeast (Figure 4B and Supplementary Figure 5B). An empty vector served as the negative control, while NubWT served as the positive control (Grefen et al. 2009). Prey and bait protein expression was verified by immunoblot analysis (Supplementary Figure 5A and 5C).

**Figure 4.**
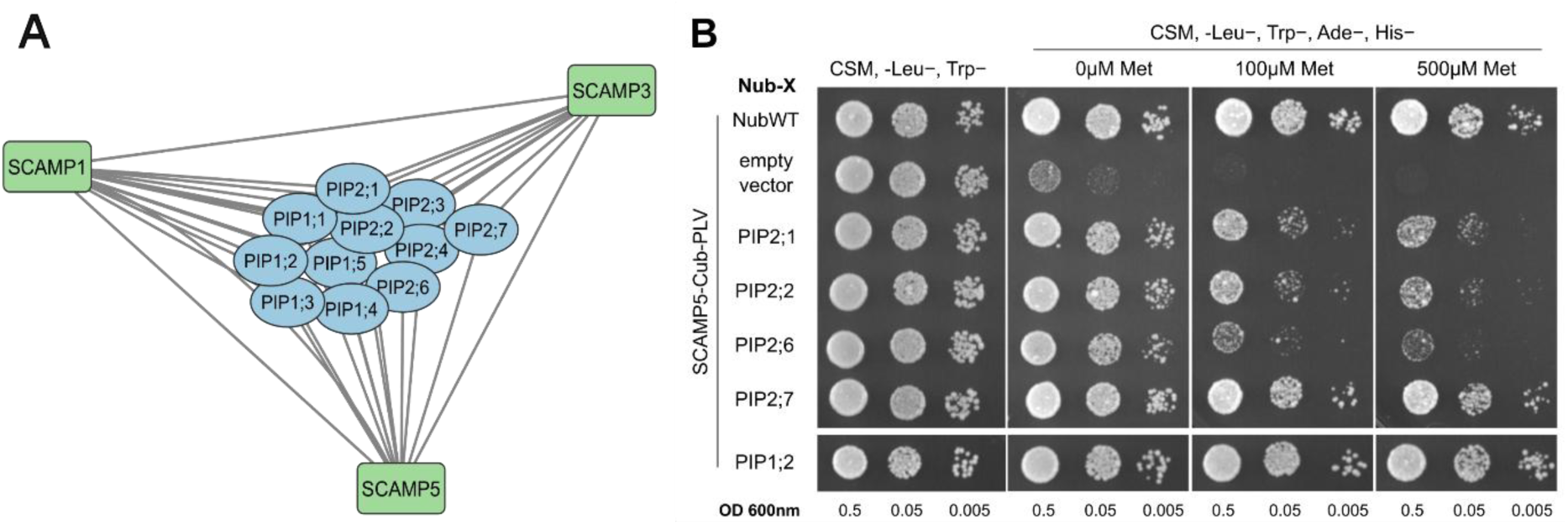
SCAMP1,3 and 5 interact with various members of the plasma membrane aquaporin family. **A)** Cytoscape-generated interaction network visualizing associations between the bait proteins SCAMP1, SCAMP3, SCAMP5 (green) and the PIP aquaporins (blue). Connections indicate statistically significant interactions identified through AP-MS analysis. **B)** Interaction analysis between SCAMP5 and several PIP2 family members as well as PIP1;2 by split ubiquitin assays. A dilution series (OD 0.5, 0.05 and 0.005) of yeast co-expressing the methionine (Met)-repressible bait construct SCAMP5-Cub-PLV and the different prey constructs (PIP1/2-NubA) were spotted onto a synthetic medium (CSM, -Leu−, Trp−) and interaction-selective medium (CSM, -Leu−, Trp−, Ade−, His−) at different dilutions with increasing Met concentrations to repress bait expression. Yeast growth was recorded after incubation for 48 h. NubWT was used as positive control and the empty vector was used as negative control. This experiment was performed once with two-independently transformed colonies for each prey-bait pair. The panel shows the results from a representative colony.

**Figure 5.**
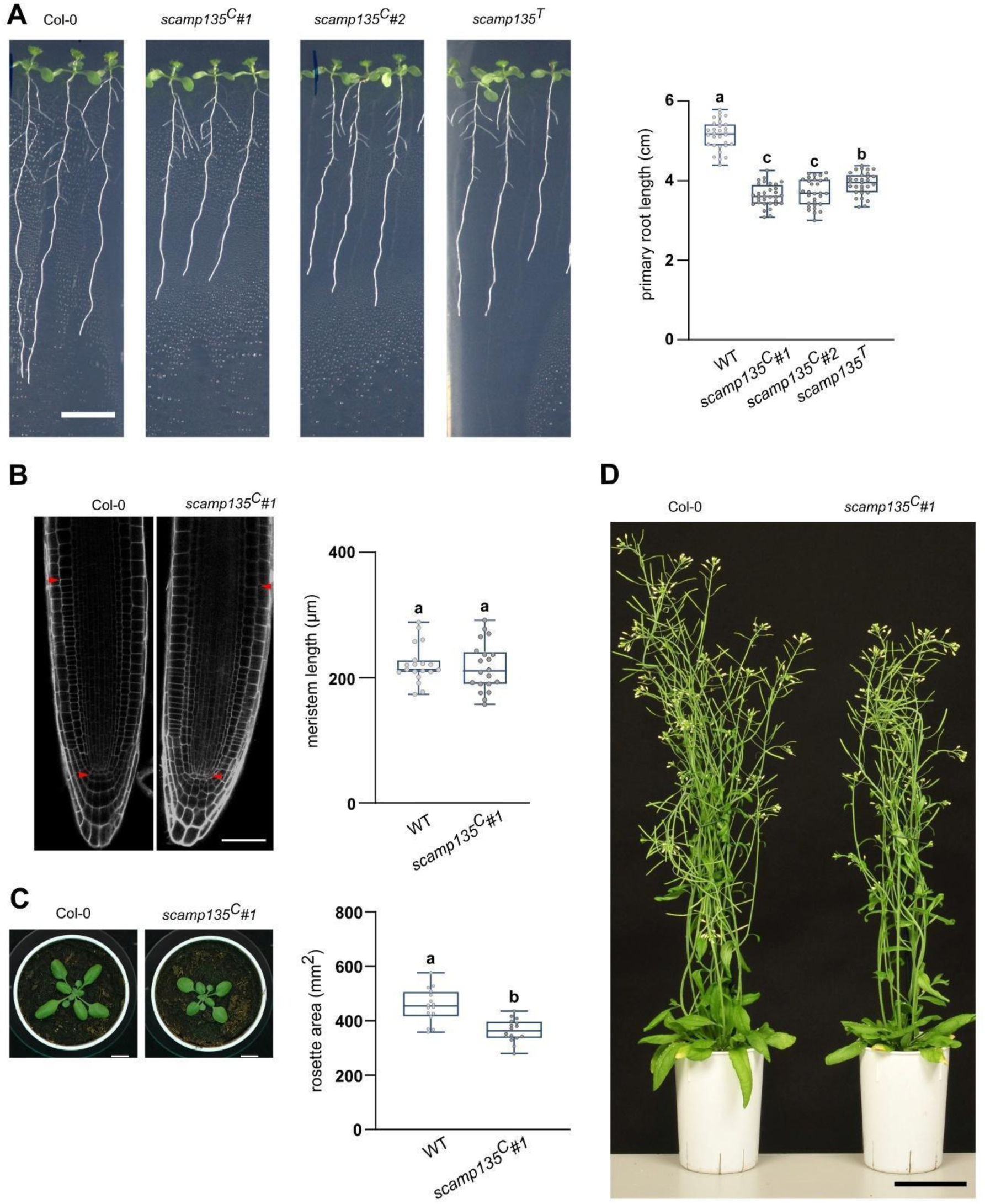
*scamp* triple mutants show mild growth defects. **A)** Representative images and box plot graph primary root length quantification of 10-day-old Col-0 and *scamp135* mutant seedlings light-grown on 1/2 MS media without sucrose. Scale bar, 1 cm. Different letters represent significant differences found by a one-way ANOVA test (P<0.05). This experiment was repeated twice. **B)** Representative root meristem images and box plot quantification of primary root meristem size of 6-day-old Col-0 and *scamp135^C^#1* seedlings stained with propidium iodide (PI). Red arrowheads mark the size of the root meristem. Scale bar, 50 µm. No significant difference was found between wild type and *scamp135^C^* mutant seedlings using a Student’s t-test (P>0,05). This experiment was repeated twice. **C)** Representative soil-grown rosette images and box plot quantification of the rosette area of 18-day-old Col-0 and *scamp135^C^* mutant plants. Different letters represent significant differences between wild type and *scamp135^C^* mutant seedlings found via Student’s t-test (P<0.0001). Scale bar, 1 cm. This experiment was repeated twice. **D)** Representative images of WT and *scamp135^C^#1* mature plants. Scale bar, 5 cm.

### *scamp135* triple mutants are affected in primary root and shoot development

To understand the physiological roles of SCAMPs in Arabidopsis, we generated mutant plants. We focused on mutants in SCAMP1, 3, and 5 based on the similarity of their overall expression pattern, their level of expression (Supplementary Figure 1) and their common interactors (Figure 4A). RT-PCR analysis confirmed the absence of *SCAMP* expression in single mutant T-DNA insertion lines (Supplementary Figure 2). However, besides mildly shorter hypocotyl lengths of *scamp1-2* compared to Col-0, no other observable aberrant phenotype was detected for the single *scamp1*, *scamp3* and *scamp5* mutants under normal growth conditions (Supplementary Figure 2).

Considering the likely functional redundancy among the SCAMPs (Figure 4A), we generated SCAMP triple knockout mutants via CRISPR/Cas9. Mutations were targeted to disrupt the localizations and functionalities of SCAMPs by targeting the first transmembrane domain. We generated two triple mutant lines, indicated as *scamp135^C^*#1 and *scamp135^C^*#2 (Supplementary Figure 6A).

Reduced end-point RT-PCR of the CRISPR-mutated *SCAMP* genes compared to the wild-type (WT), presumably caused by nonsense-mediated mRNA decay, confirmed the disruption of *SCAMP1*, *3* and *5* (Garneau et al. 2007; Lykke-Andersen et al. 2015) (Supplementary Figure 6B). Macroscopically, the triple mutants exhibited shorter roots, a smaller rosette area compared to WT at early developmental stage and a slightly reduced stature at maturity (Figure 5A, 5C and 5D). The shorter root phenotype was phenocopied in a *scamp135* mutant line derived from crossing the single T-DNA insertion mutants (*scamp135^T^*) and in independent alleles of *scamp12345^C^* quintuple mutants (Supplementary Figure 6E). The fact that the quintuple mutant phenotype resembles the triple mutant points to SCAMP1, 3 and 5 as the major isoforms functioning in roots, which is in agreement with their expression pattern (Supplementary Figure 1). Root meristem lengths were comparable between *scamp135^C^#1* and WT, pointing to cell elongation as a major cause for the observed reduction in main root length (Figure 5B), which would be in line with the observed interaction between SCAMPs and PIPs (Figure 4) and a role for PIPs in cell elongation (Hachez et al. 2006; Fricke and Knipfer 2017; Israel et al. 2022). Introducing SCAMP5-GFP into the *scamp135^C^#1* mutant background partially rescued the root length phenotype to that of the *scamp13^C^* double mutant (Supplementary Figure 6F). Considering SCAMPs’ presumed role in trafficking, we tested if secretion was affected. We could not detect a difference in secretory capacity between the TGN/EE and the cell plate, but we could identify a mild, yet significant, decrease in mucilage secretion upon seed imbibition when *scamp* triple and quintuple mutants were compared with WT. The observed decrease in secretion was however much smaller than the defect that is present in the *picalm1a/b* mutant, which served as a positive control for this assay (Fujimoto et al. 2020) (Supplementary Figure 6C and 6D).

**Figure 6.**
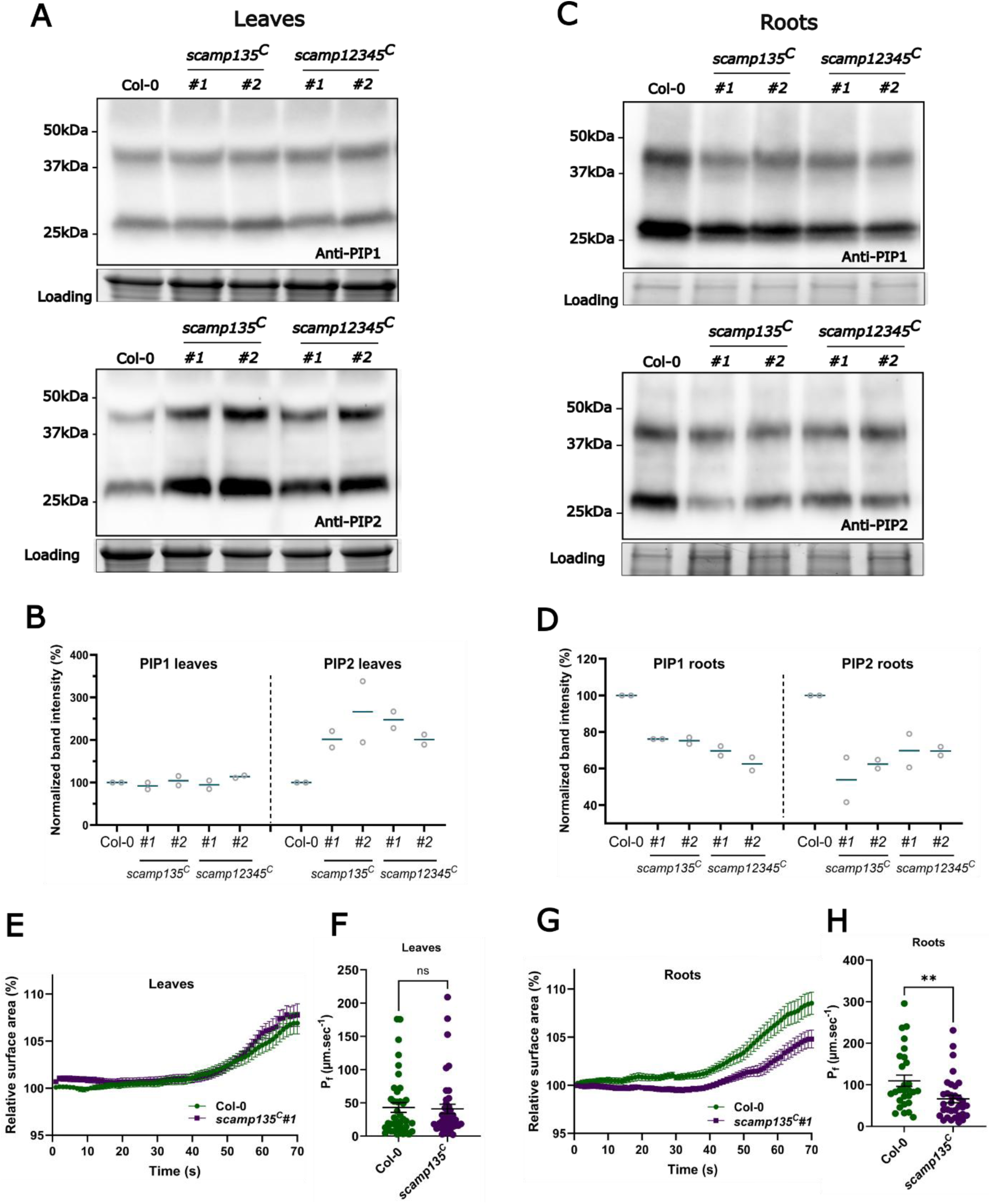
PIP accumulation and water transport are reduced in *scamp* mutants. **(A and C)** Western blot analysis of WT Col-0 and two independent lines *(#1* and *#2*) from both *scamp135^C^* and *scamp12345^C^* mutants using antibodies against endogenous PIP1 and PIP2 in leaves (A) and in roots (C) from 7-day-old seedlings. The stain-free gel serves as a loading control. **B and D)** Dot graph representing relative band intensity normalized to the loading control (%) of PIP1 and PIP2 monomer and dimer in leaves (B) and roots (D). The dots represent the quantification of two independent experiments and the line presents the mean value of both measurements. PIP1 and PIP2 monomer and dimer proteins have theoretical molecular weights of ∼30 kDa and ∼55 kDa respectively. Note that in our experimental conditions, PIPs display an apparent molecular weight below the expected one. Representative immunoblots are shown. Blots of the other experiments can be found in the source file. (**E and G)** Root and leaf protoplast swelling time course (70 sec) showing the relative surface area of WT and *scamp135* triple mutants upon exposure to a hypotonic challenge (values represent mean ± SE). **(F and H)** Osmotic water permeability values (P_f_) (mean ± SE) based on the graphs in E and G. ns and double asterisks indicate non-significant and significant (Student’s *t*-test, p = 0.01) differences from the control, respectively. Data were obtained from two independent experiments for both root and leaf protoplasts, with a total of 28-42 protoplasts for each genotype.

### The abundance of PIP1 and PIP2 is altered in *scamp* mutants

To identify whether the SCAMPs control aquaporin trafficking, we compared PIP levels between wild type plants and triple and quintuple *scamp* mutants. PIP1 and PIP2 abundance was consistently decreased in roots in all *scamp* mutant lines compared to the wild type. In contrast, in leaves, PIP2 levels were increased in *scamp* mutants. A similar trend, although less pronounced and more variable between the different lines could be observed for PIP1 levels (Figure 6A-D). Taken together, the data indicate that SCAMPs control the abundance of various PIP isoforms in an organ-dependent way. To assess whether the reduced PIP levels correspond to reduced PIP levels at the PM, we performed protoplast swelling assays (Moshelion et al., 2004; Shatil-Cohen et al., 2014). Independent experiments showed significantly reduced water transport in *scamp135* mutant root protoplasts compared to wild type under hypotonic conditions. No significant changes were observed for leaf protoplasts (Figure 6E-H). Our protoplast swelling results are in agreement with our Western blotting results and show that PIP abundance at the PM is specifically reduced in *scamp135* mutant roots.

### *scamp* mutants are less sensitive to drought-induced leaf wilting

Considering the pivotal roles of PIPs in water transport and drought responses (Chen et al. 2021; Liu et al. 2023), we examined the effects of drought on our *scamp* mutants. We first subjected Col-0 and scamp135C#1 mutants to mannitol as a proxy for mild to moderate drought stress. Whereas *scamp135^C^* mutants displayed shorter roots compared to the WT under standard growth conditions, root growth of the *scamp* triple mutants was less affected when grown on mannitol, leading to a higher normalized root growth when compared to WT seedlings (Supplemental Figure 7A).

Subsequently, single and double mutants derived from T-DNA insertion lines, as well as the *scamp135^C^#1* triple mutants were analysed using an automated weighing, imaging, and watering machine (WIWAM) system (Skirycz et al. 2011). Arabidopsis plants were grown for 2 weeks under well-watered conditions. After two weeks, half of the plants continued to receive water while watering was stopped for the other half. All plants were imaged for another 10 days. The rosette area of the plants that did not receive any water steadily increased until 7 days after watering was stopped (7 DAWS). We compared this peak rosette area (7 DAWS) with the rosette area at 10 days after watering stopped (10 DAWS) (Figure 7A). Rosette area ratios (10 DAWS / 7 DAWS) of the single, double, and triple *scamp* mutants were all significantly larger than those of Col-0, indicating that loss of SCAMP function makes Arabidopsis plants less sensitive to drought-induced leaf wilting (Figure 7B).

**Figure 7.**
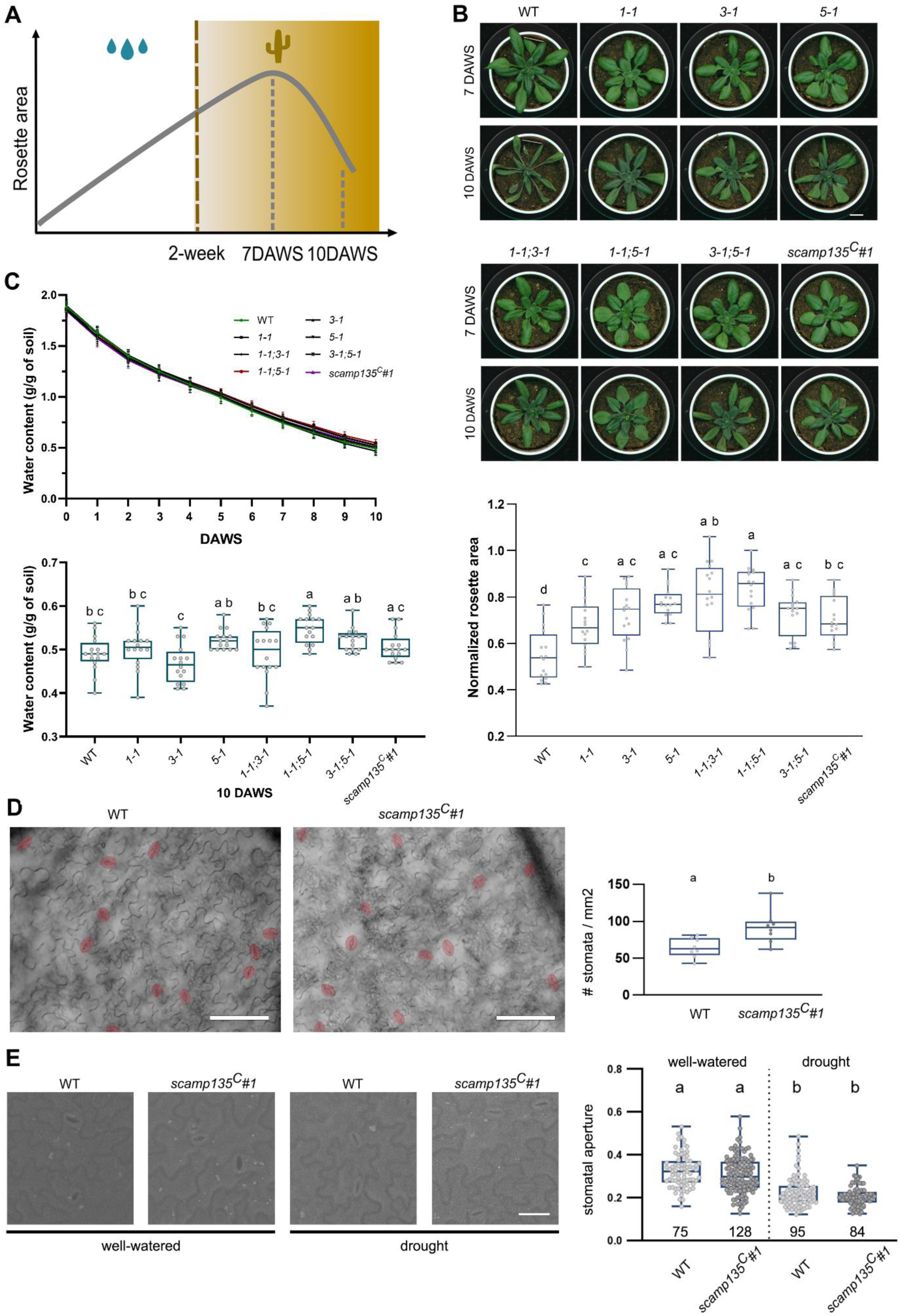
*scamp* mutants show reduced drought sensitivity compared to WT. **A and B)** Schematic representation (A) and representative rosette images and quantification (B) of the WIWAM drought sensitivity experiment. Arabidopsis plants were grown under well-watered conditions for 2 weeks and then watering was stopped (dashed line) and plants were followed for an additional 10 days (10 DAWS). The rosette area gradually increased and reached a maximum 7 days after watering was stopped (7 DAWS). The quantification represents the ratio of the rosette area at 10 DAWS over the maximum rosette area at 7 DAWS for the different genotypes that were tested. Scale bar, 1 cm. **C)** Changes in soil water content following watering stop of WT and *scamp* mutants. Water soil content (g/g) was calculated and followed from 16 DAS (0 DAWS) until 26 DAS (10 DAWS). The quantification represents water soil content values at 10 DAWS. Different letters represent significant differences evaluated by a one-way ANOVA analysis including a Tukey’s multiple comparison test (0.0001<P<0.02). **D)** Representative images with indication of mature stomates (red) and quantification of stomatal density on the abaxial side of the third leaf of three-week-old soil-grown WT and *scamp135^C^#1* mutant plants. Four to six non-overlapping images were used for the quantification. For each genotype, eight leaves from different plants were measured. Each dot represents the average of the four to six images from one biological replicate. Scale bar, 100 µm. Different letters represent significant differences evaluated by a two-tailed unpaired t test (0.001<p<0.01). **E)** Stomatal aperture under well-watered and drought conditions of soil-grown plants. Representative dental resin imprint SEM images and quantification of stomatal aperture (width over length) of the third leaf epidermis of three-week-old WT and *scamp135^C^#1* mutants. The values under the box plots (n) indicate the number of stomata that were measured from individual third leaves from independent Arabidopsis Col-0 (5 and 5 plants) and *scamp135^C^#1* (4 and 6 plants) with or without drought, respectively. Different letters represent significant differences evaluated by a Kruskal-Wallis test including a Dunn’s multiple comparison test (p<0.0001). Scale bar, 30 µm.

Wilting tolerance can be caused by changes in roots and leaves (Kuromori et al. 2022). Soil water content over the drought treatment was similar between all the genotypes, with only the *scamp1-1/1-5* mutant showing a statistical significant difference compared to the wild type control plants (Figure 7C). Triple mutant *scamp135^C^#1* soil-grown plants had a slightly higher stomatal density than WT plants under well-watered conditions (Figure 7D) and detailed analysis of stomatal dynamics under well-watered and drought conditions did not reveal differences between WT and the *scamp135^C^#1* soil-grown plants (Figure 7E). These results indicate that the reduced wilting phenotype of the *scamp* triple mutants cannot be easily explained by reduced water loss due to altered stomatal dynamics or densities, nor by differential soil water availability despite that *scamp* mutants have slightly smaller rosette areas (Supplementary Figure 7B and 7C).

Osmotic stress causes removal of PIPs from the PM (Hachez et al., 2014; Pou et al., 2016). To analyse how PIP abundance would change upon drought, we mimicked drought stress by sorbitol and monitored PIP levels in wild type and *scamp135* triple mutant plants. Sorbitol treatment caused a strong reduction of PIP1 and PIP2 levels in wild type roots, whereas in *scamp135* mutant roots, the reduction was less pronounced. In contrast, in leaves, similar reductions in PIP1 and PIP2 levels were observed between both backgrounds (Figure 8A-D). We therefore conclude that the low steady-state levels of PIPs in the *scamp135* mutant roots may act as a priming mechanism, allowing the plants to respond better to conditions of reduced water availability.

**Figure 8.**
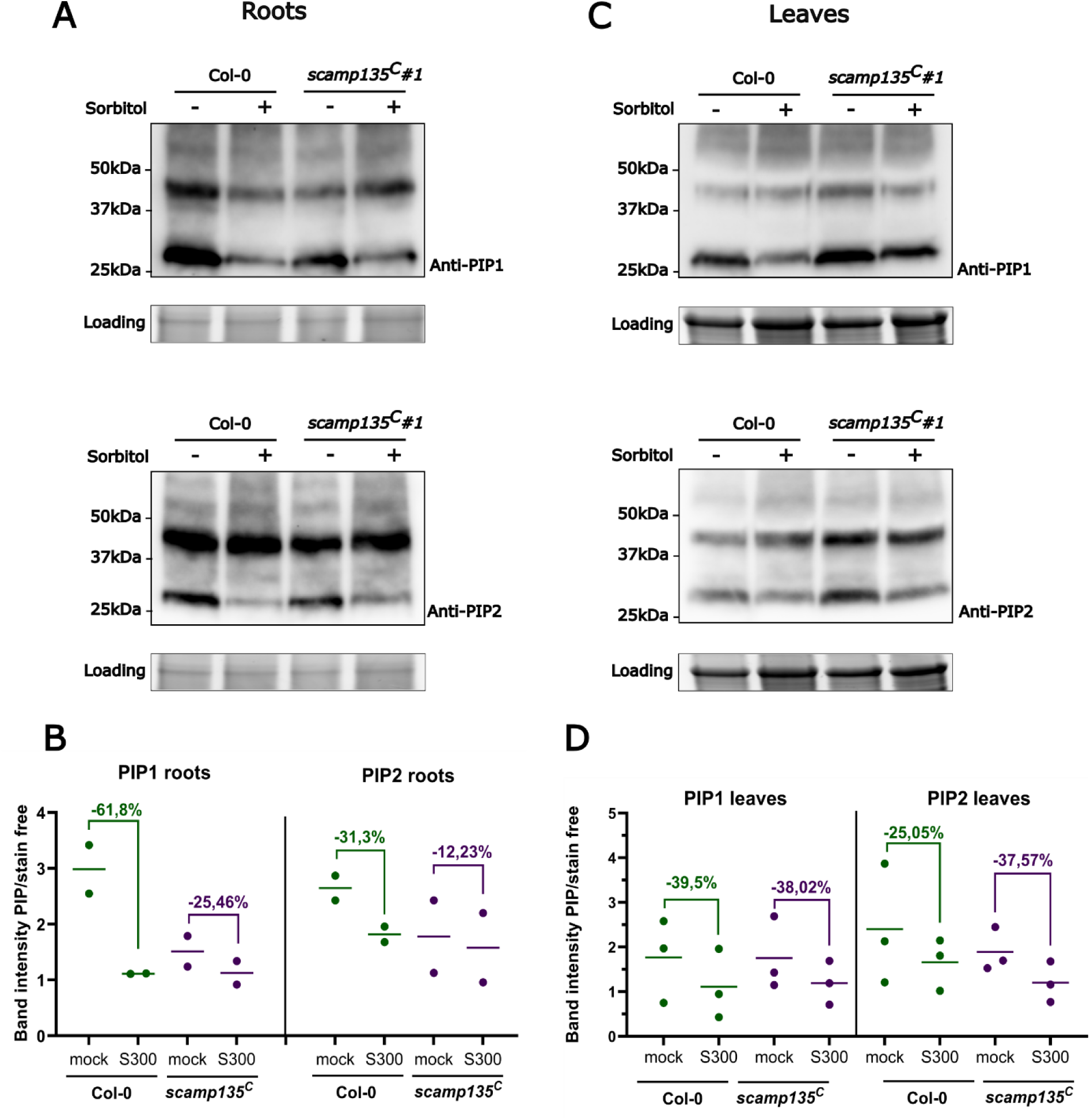
Sorbitol treatment reduces PIP levels differentially between wild-type and *scamp* mutants. **A and C)** Western blot analysis of WT Col-0 and *scamp135^C^#1* using antibodies against endogenous PIP1 and PIP2 in roots (A) and in leaves (C) from 7-day-old seedlings treated with (+) or without (-) 300mM sorbitol treatment. The stain-free gel serves as a loading control. PIP1 and PIP2 monomer and dimer proteins have theoretical molecular weights of ∼30 kDa and ∼55 kDa respectively. Note that in our experimental conditions, PIPs display an apparent molecular weight below the expected one. Representative immunoblots are shown. Blots of the other experiments can be found in the source file. **B and D)** Dot graph representing band intensities normalized to the loading control of PIP1 and PIP2 monomer and dimer in roots (B) and leaves (D). The dots represent the quantification of independent experiments and the line presents the mean value of both measurements. The percentage of PIP degradation under sorbitol (S300) treatment relative to mock conditions for each genotype is shown.

## Discussion

### SCAMP internalization depends on a linear N-terminal motif

The SCAMP protein family has been extensively studied in animal cells (Hubbard et al. 2000; Castle and Castle 2005; Lin et al. 2005; Müller et al. 2006; Aoh et al. 2009; Diering et al. 2009; Fjorback et al. 2011; Zaarour et al. 2011; Lee et al. 2021). In contrast, our understanding of the role of these proteins in plants is very fragmented, with only few studies using them merely as markers (de Keijzer et al. 2017; Yamada et al. 2025) or addressing their trafficking in heterologous systems like tobacco BY-2 cells (Sheung et al. 2007, 2008). Functionally, the limited literature describes very diverse roles for plant SCAMPs, ranging from salt stress tolerance in soybean and cotton to restricting bark deposition in poplar and from pollen tube elongation in lily to a role in stomatal response to ABA in Arabidopsis cotyledons (Wang et al. 2010; Obudulu et al. 2018). Here, we characterized the trafficking and the role of this protein family in the model plant Arabidopsis using live cell imaging, interactomics and higher order mutant analysis. The Arabidopsis genome encodes five SCAMPs. SCAMP1, 3 and 5 seem to be the major isoforms expressed in root cells. These three SCAMPs appear to localize to the PM and to endosomes. The endosomal pool of SCAMP5 consisted mainly of TGN/EE. This is in agreement with what is reported for PpSCAMP4 (de Keijzer et al. 2017). Internalization of SCAMP5 and SCAMP3 from the PM relies on N-terminal NPF motifs, which were shown to bind directly to the first EH domain of the AtEH1/Pan1 subunit of the endocytic TPLATE complex (TPC) (Gadeyne et al. 2014; Yperman et al. 2021). The slightly dominant negative effect of SCAMP5_mNPF on root growth indicates that SCAMP5 likely has a function in endocytosis beyond merely being a cargo that has to be retrieved from the PM. The recognition of SCAMP5 by the TPC strongly hints at TPC-dependent internalization of SCAMP5. Given the general role of TPC in endocytosis, this was not directly tested here as this would require establishing a complemented *eh12* double mutant line containing a functional TPC lacking the ability to recognize SCAMP5 by mutating W49 and W51 in the first EH domain of AtEH1/Pan1 and AtEH2/Pan1 respectively (Yperman et al. 2021).

The evidence for the importance of ubiquitination as a mechanism for cargo recognition for endocytic internalization in plants largely exceeds that of the importance of the short linear motifs (Arora and van Damme 2021; De Meyer et al. 2023; Saeed et al. 2023). SCAMP5 can be ubiquitinated at two residues in its N-terminal domain (Grubb et al. 2021). Although the correlation between the presence of the double NPF motif and the possibility for SCAMP5 to be ubiquitinated was not tested directly, SCAMP5 lacking both NPF motifs was hardly internalized, indicating a major role for this motif in its recognition as endocytic cargo.

### SCAMPs function as dimers

Many of our results argue for SCAMPs to operate as dimers. SCAMPs were found as top interactors in our interactomics experiments and we confirmed the homo- and hetero-dimerization capacity of SCAMPs via their N-terminal alpha-helical domain. The most likely explanation for SCAMP5_mNPF to act in a dominant-negative manner to disrupt SCAMP5-GFP localization is that SCAMPs also internalize as dimers. In agreement with this, we found evidence for hetero-dimerization in the PM and at the TGN. In animal cells, truncated SCAMP2 also redistributed wild-type SCAMP2, which is consistent with our findings on SCAMP dimerization (Lin et al. 2005).

### SCAMPs are not essential for Arabidopsis development under standard conditions

Single knock-out mutants in *SCAMPs* do not show obvious developmental defects. Higher-order mutants have a slightly decreased seedling root growth, likely due to perturbed cell elongation. This observation hints at functional redundancy between SCAMPs. The quintuple mutants had no aggravated effect on root growth compared to the triple mutants, indicating that SCAMP2 and 4 do not play a major role in root development. Also, the viability of the *scamp* quintuple mutant shows that these proteins have no essential function under standard greenhouse conditions or that parallel pathways are operational. In contrast, silencing two SCAMPs in poplar led to increased deposition of saccharides and phenolic components in the stem, suggesting that *Populus* SCAMPs act as a safeguard to suppress the secretion of cell wall precursors (Obudulu et al. 2018). Given the observed localization pattern of SCAMPs in Arabidopsis roots, a role for SCAMP1, 3 and 5 in retrograde or anterograde post Golgi transport is indeed expected. However, in contrast to what is reported in poplar, our results in Arabidopsis rather favour a positive role in secretion for SCAMPs as seed mucilage secretion in *scamp* triple and quintuple mutants was slightly reduced compared to that of WT. We did not test bark accumulation in the stem as Arabidopsis, being an annual herbaceous plant, accumulates limited woody tissue in its stem (Grennan 2006; Melzer et al. 2008).

### PIP abundance is altered in higher order *scamp* mutants

PIP aquaporins are the most prominent protein family in our interactomics analysis from Arabidopsis seedlings expressing SCAMP1, SCAMP3 and SCAMP5. Yeast assays confirmed the interaction between SCAMP5 and both type 1 and type 2 PIPs to be direct. Higher order Arabidopsis *scamp* mutants have clearly and robustly affected PIP1 and PIP2 levels and we furthermore observe clear differences in PIP levels between above- and belowground organs.

Three water-transporting pathways move water through the whole plant, the apoplastic path through cell walls, the symplastic path going through plasmodesmata, and the transcellular path where the water crosses membranes. The latter path is controlled by PIPs (Javot and Maurel 2002). Being PM aquaporins, the direct and positive roles of PIPs in hydraulic conductivity are well-studied. Aquaporins are involved in cell expansion and division in distinct tissues. For example, the expression of several PIPs in maize primary roots precedes the onset of cell elongation (Hachez et al. 2006). Hence, it was suggested that the synthesis and accumulation of aquaporins increases the hydraulic conductivity of the membranes, which is required for cell elongation and organ development. In line with this, the shorter primary root length, without a major effect on meristem size in the *scamp135^C^* mutants, suggests reduced cell elongation. Whether this is caused by reduced water transport capacity or by altered proton pump ATPase activity remains to be determined as our interactomics analysis also revealed several AHA proteins as common interactors of the three SCAMPs (Supplemental Figure 4). The smaller rosette area in *scamp135^C^* is also not that easily explained by reduced hydraulic conductivity of the cell PM as PIP levels are not reduced in *scamp135* mutant leaves.

PIPs are highly regulated water channels that contribute to drought tolerance and the maintenance of water homeostasis (Chaumont and Tyerman 2014). In Arabidopsis, *pip2;8* single and *pip2;1 pip2;2* double mutants show drought tolerance, while plants overexpressing PIP2;1 and PIP2;8 are more sensitive to drought than the WT (Chen et al. 2021; Liu et al. 2023). The stomatal aperture is smaller in *pip2;8* plants, and ABA-induced stomatal closure is enhanced in *pip2;1 pip2;2* mutants, suggesting that reducing the amount/activity of these PIPs in the PM increases drought tolerance through stomatal closure (Liu et al. 2023). PIPs indirectly also controls many aspects of plant development as quintuple *pip1* mutants are affected in various aspects of plant development(Wang et al. 2021).

PIPs also play important roles in water uptake at the root level and during water stress. Arabidopsis *pip2;2* mutants exhibited significant decrease in the hydraulic conductivity in cortex cells and in whole roots (Javot et al. 2003), and water stress can modify the root hydraulic conductivity (Chaumont and Tyerman 2014). In addition, inhibiting PM targeting of PIPs also affects the cell and root water membrane permeability. Mutation of ER exit motif found in PIP2;1 resulted in its ER retention and a decrease in Arabidopsis root hydraulic conductivity (Sorieul et al. 2011).

Our results in Arabidopsis support a complex role for the SCAMPs in the trafficking of the plasma membrane-localized aquaporins. We observed reduced water transport under hypo-osmotic conditions, caused by reduced abundance of PIPs at the PM in roots, but not in leaves, which might be related to the low expression of *SCAMPs* in leaf mesophyll cells (Supplementary Figure 1). Whether SCAMPs, next to their abundance, also modulate the water transport activity of the PIPs remains to be determined.

### Impairing Arabidopsis SCAMPs reduces leaf wilting under drought conditions

Single mutants in *SCAMP1*, *3* and *5* all presented a drought tolerance phenotype at the level of the rosette area and this tolerance was similar to that of the double and triple mutant combinations. This similar phenotype for single and higher order mutant combinations aligns with the importance of heterodimerization for SCAMP function.

Previously, single Arabidopsis *scamp* mutants were reported to show resistance to ABA-induced stomatal closure in isolated cotyledons (Bourdais et al. 2019). Reduced wilting, which we observe for our single and higher order *scamp* mutants, is intuitively not easily linked to open stomata. Our results, using leaves of soil-grown plants instead of cotyledons show no major difference in stomatal opening between WT and *scamp* higher order mutants both under well-watered conditions as well as under drought treatments. Whether the observed discrepancy in stomatal dynamics is due to the tissue or the treatment remains unclear at the moment. Our results however suggest that the observed drought tolerant phenotype of the *scamp135^C^* triple mutants cannot be explained by stomatal aperture defects or by reduced stomatal density. The reason for the reduced wilting of the *scamp* mutants is therefore likely associated with increased water retention by the roots during drought rather than by differential water loss via the leaves. The *scamp* mutant plants are slightly smaller than WT when grown under well-watered conditions, yet the soil water content upon drought was equal indicating that the observed rosette fitness is not caused by drought avoidance.

Functionally, our results are in agreement with previous reports showing increased drought tolerance by reducing PIP plasma membrane levels via Endoplasmic reticulum-associated PIP degradation (Lee et al. 2009; Chen et al. 2021). Whether the reduced PIP abundance in *scamp135* mutants is caused by the activation of ER-dependent PIP degradation remains to be determined. The causality of the drought tolerance of the *scamp* mutants might also be in part inferred by functions of SCAMP that are independent of their interaction with the PIPs. In addition to the size of the leaves, other factors like leaf angle, leaf thickness, and cuticle deposition as well as root architecture play significant roles in influencing a plant’s response to drought conditions (Smith and De Smet 2012; Yavas et al. 2024).

## Methods

### Molecular cloning

Genomic SCAMP fusions and motif-mutated constructs were cloned via Gateway recombination. An upstream region (2.5 kb) before the start codon of each SCAMP (pSCAMP) was cloned into pDONRP4P1R and a genomic SCAMP sequence without a stop codon was cloned into pDONR221. Motif-mutated SCAMP3 or SCAMP5 isoforms were generated by mutagenesis PCR from the pDONR221 containing full-length SCAMP3 or SCAMP5 plasmids by combining sewing and mutation primers and cloning the PCR product into pDONR221. All primers are listed in Supplementary Table 1.

pSCAMP::SCAMPs/SCAMP3_mHSF/SCAMP5_mNPF/SCAMP5_mY-eGFP were made by combining pDONRP4P1R-pSCAMP, pDONR221-SCAMP/SCAMP3_mHSF/SCAMP5_mNPF/SCAMP5_mY, and pDONRP2RP3_mGFP (Karimi et al. 2007) in the pB7m34GW or pH7m34GW backbone (Karimi et al. 2005). pSCAMP5::SCAMP5_mNPF-mCherry was obtained by combining the generated entry clones, and pDONRP2RP3_mCherry (Mylle et al. 2013) in the pH7m34GW backbone (Karimi et al. 2005). All constructs used are listed in Supplementary Table 1.

Constructs used in the rBiFC were built using the 2 in 1 BiFC system (Grefen and Blatt 2012). Genomic SCAMP5 and SCAMP3 without a stop codon were cloned into pDONR221P3P2 and pDONR221P1P4, respectively. Genomic BRI1, without a stop codon, was amplified and cloned into pDONR221P1P4. pDONR221P3P2_SCAMP5 was combined with pDONR221P1P4_SCAMP5 (Arora et al. 2020), pDONR221P1P4_SCAMP3, and pDONR221P1P4_BRI1 in the pBiFct-2in1-CC empty backbone vector to generate the destination vectors.

For Y2H, the coding sequence of all SCAMP N-terminal fragments was amplified with primers having a SmaI enzyme digest site as a hangover. pGBKT7 and pGADT7 were linearized with SmaI, then combined with the fragments via NEBuilder. The final vectors were checked by sequencing.

For SUS, cDNAs of the gene of interest were PCR amplified using the corresponding attB1 and attB2 primers (Supplementary Table 1) and cloned into pDONR221. The entry clones were recombined with the SUS destination vectors pMetYC-DEST, pNX35-DEST, and pXNubA22-DEST (Grefen et al. 2009). pDONRP1P2_PIP2;7 and pMetYC-PIP2;7 were already published (Hachez et al. 2014). pMetYC-DEST was used to produce the Met-repressible bait construct SCAMP5-Cub-PLV and PIP2;1/2/6/7-Cub-PLV. pNX35-DEST was used to produce the prey construct NubG-SCAMP5. pXNubA22-DEST was used to produce the prey constructs PIP2s-NubA and PIP1;2-NubA. The NubWT positive control fragment was obtained from the pNubWT-Xgate vector (Grefen et al. 2009).

The plasmids used to create the CRISPR mutants, the protocol and backbone vector (see Supplementary Table 1) were used as previously described (Decaestecker et al. 2019; Develtere et al. 2024). Briefly, four different guide RNAs (gRNAs) for each *SCAMP* were designed by CRISPOR (http://crispor.tefor.net/). Two destination vectors were made using the Green Gate cloning system (Lampropoulos et al. 2013) and each vector contained two gRNAs for each *SCAMP*, which means that 6 gRNAs for three genes were loaded in one expression vector.

All entry clones were verified by Sanger sequencing. All destination vectors were checked by sequencing the borders of the backbone plasmids.

### Plant material

The Arabidopsis plants used for this study are in the Columbia-0 ecotype. Flowering Arabidopsis plants were dipped with *Agrobacterium tumefaciens* C58C1 containing various plasmids in 10% sucrose buffer with 0.1% Silwet L-77 (LEHLE SEEDS, Cat. No. VIS-30) (Clough and Bent 1998). Primary generation (T1) Arabidopsis transgenic plants were selected with antibiotics and were propagated. All transgenic lines are listed in Supplementary Table 2.

The *scamp* single and triple homozygous T-DNA insertion lines were confirmed by genotyping PCR. The lines and the primers used are listed in Supplementary Table 1. The CRISPR mutants were generated via the fluorescence-accumulating seed technology (FAST) system (Shimada et al. 2010; Decaestecker et al. 2019). T2 seeds were screened for the absence of fluorescence to discard the CRISPR editing construct, and T2 plants were sequenced to check for gene editing. Homozygous mutants were verified by sequencing.

The lines combining SCAMP5 with the ST-mRFP (Renna et al. 2005), VHA-a1-mRFP (Fecht-Bartenbach et al. 2007) and mRFP-ARA7 (Ueda et al. 2004) markers were generated by dipping the genomic SCAMP5 fusion construct with GFP into the marker lines.

The lines expressing SCAMP1-GFP and SCAMP5-mCherry were generated by dipping the SCAMP5-mCherry construct into the line expressing SCAMP1-GFP. The lines expressing SCAMP5-GFP and SCAMP5_mNPF-mCherry were generated by dipping the SCAMP5_mNPF-mCherry construct into the line expressing SCAMP5-GFP. The *scamp135^C^*#*1* line was transformed with pSCAMP5::SCAMP5-GFP to address the mutant complementation. The *scamp13^C^*#1 line was generated by back-crossing *scamp135^C^*#1 with wild-type, growing the heterozygous plants and selecting the double mutant in the offspring generation.

T3 Arabidopsis plants homozygous for the construct and the T-DNA mutation, *pSCAMP5::SCAMP5-eGFP*/*scamp5-1(-/-)*, *pSCAMP5::scamp5_mNPF-eGFP*/*scamp5-1(-/-)*, and *pSCAMP5::scamp5_mY123-eGFP*/*scamp5-1(-/-)* were used for imaging and phenotyping by comparing them to *scamp5-1(-/-)*. Two independent transgenic lines were selected based on fluorescence intensity.

### Phenotypical analysis

Unless otherwise specified, seeds were sterilized by chlorine gas and then were sown on 1/2 MS medium plates without sucrose. The plates were incubated in a growth chamber under continuous light conditions at 21 °C after 2 days of stratification at 4 °C.

To compare the primary root (on 1/2MS and on mannitol containing plates), and hypocotyl length, 10 to 13-day-old light-grown seedlings and 4-day-old Arabidopsis seedlings that were grown in the dark were used respectively. Measurements were done using the NeuronJ plugin (Meijering et al. 2004) in the ImageJ software package (https://imagej.net/ij/).

To assess the rosette area of the *scamp135^C^* mutants in Figure 7 and the drought tolerance of the various *scamp* mutants, plants were sown directly in soil and stratified for two days at 4 °C. Pots were transferred on the WIWAM system and growth was monitored (Skirycz et al. 2011). Arabidopsis plants were grown at 21°C using a 16 hours day/8 hours night cycling environment. Arabidopsis plants were imaged every day. Watering was stopped after 2 weeks (16 days after sowing, DAS) for half of the plants, while the other half of the plants continued to grow under well-watered conditions. Rosette sizes were automatically calculated by the WIWAM2 system (Skirycz et al. 2011). This yielded initial growth curves under well-watered and under drought conditions, allowing us to determine the rosette sizes of two-week old plants (16DAS, Supplementary Figure 7B) as well as the maximal rosette size of the plants after watering was stopped (7 DAWS, 23 DAS). We also calculated the rosette size of the different genotypes under well-watered conditions at the 23 DAS timepoint (Supplementary Figure 7C).

For the rosette area quantification in Figure 7, 23 DAS (7DAWS) and 26 DAS (10 DAWS) plants are presented and quantified. The drying rosette areas were not adequately quantified by the WIWAM software package. Therefore, rosette areas of all genotypes at 7 DAWS and 10 DAWS were measured manually using the polygon selection tool in ImageJ. Representative rosette pictures of plants are presented in Figure 7.

Soil water content was calculated using the formula:

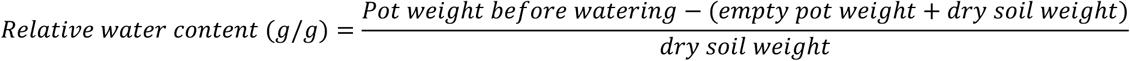

This yielded the water retained per gram of oven-dried soil.

Visualization of mucilage secretion in seeds was managed by ruthenium red staining. After hydration for 2 hours in Tris 10 mM pH 7.5, seeds were stained with 0.1% (w/v) ruthenium red (Sigma Aldrich) for 1 hour. Then, they were rinsed twice in water before imaging using a Leica M165 FC stereomicroscope. For each line, about 40 individual seeds were analysed using ImageJ. For quantifying the size of the seeds, a threshold range from 50 till 255 was used and for the seed + mucilage area, a range from 200 till 255 was applied and the image was inverted before quantifying the area using the particle analysis tool in ImageJ.

### RT-PCR

RT-PCR to confirm the homozygosity of the T-DNA lines and to check nonsense mRNA decay in the *scamp* CRISPR mutants was done as follows: Seedlings were harvested after 10 days on 1/2 MS at CL conditions. Total RNA of mutants and WT was extracted with the ReliaPrep™ RNA Miniprep System (Promega, Z6010). cDNA was made from 1 ug mRNA using the RevertAid First Strand cDNA Synthesis Kit (Thermo Scientific) or with the qScript Ultra SuperMix (Quantabio, 95217). PCR was performed on 2 ul the cDNA using GoTaq Flexi DNA Polymerase (Promega) in a total volume of 50 ul. Primers at the 5’ and 3’ UTR region of *SCAMPs* were chosen to amplify *SCAMPs*. *ACTIN2* expression was used for normalization. All primers are listed in Supplementary Table 1. For the quantification in Supplementary Figure 5B, bands from three gels were quantified using Image Lab 6.0 (Bio-Rad).

### Tobacco infiltration

3-week-old *Nicotiana benthamiana* plants grown in the greenhouse under long-day conditions were used for leaf infiltration (Sparkes et al. 2006) for the rBiFC experiments. Transformed Agrobacteria strains were incubated in liquid YEB medium for 3 days. The pellets were washed in an infiltration buffer (10 mM MgCl2, 10 mM MES pH 5.7), then resuspended in an infiltration buffer with 100 μM acetosyringone. The OD600 value was adjusted to 1.0. The cultures were incubated on a rotating wheel for 2-3 hours at room temperature before injecting the tobacco lower epidermis leaves.

### Live cell imaging

#### Transient expression analysis in *N. benthamiana*

1. *N. benthamiana* leaves were imaged two days after infiltration. All standard imaging was done using an inverted Leica SP8X confocal microscope equipped with a 40x (NA 1.10) or a 25x (NA 0.95) water immersion corrected objective using white-light laser excitation. eGFP (excitation 488 nm, emission 500-550 nm), YFP for rBiFC (excitation 514 nm, emission 520-550 nm), and mCherry/mRFP/PI (excitation 561 nm or 587 nm, emission 600-650 nm or 650-750 nm) were visualized on Hybrid detectors and fluorescence was collected with a time-gating window between 0.3-6.0 ns. Images of the dual-colour lines were acquired in line sequential mode. Samples were mounted between slide and coverslip.

#### FRET-FLIM interaction analysis

For FRET-FLIM, images were collected using the Falcon (Fast Lifetime CONtrast) module of a Leica Stellaris 8 confocal, equipped with an HC PL APO CS2 60x water immersion objective (NA 1.2). The donor lifetime values were determined by time-correlated single-photon counting (TCSPC) in Arabidopsis root cells expressing the proteins of interest fused to either eGFP (donor) or mCherry (acceptor). The 488 laser of the Stellaris 8 white-light laser, used at a frequency of 40 MHz, was used for donor fluorescence excitation. In order to prevent pulse pile-up, the laser power was adapted to a maximum count rate of one photon per laser, nicely visualized by a vertical red line in the pixel intensity histogram. Image acquisition was done at zoom 3 with a resolution of 512 x 512 pixels with a pixel size of 0.120 um/pixel, and a pixel dwell time of 3,162 us. Photons were accumulated during 12-line repetition scans to obtain appropriate photon numbers (around 1000 peak counts) for reliable statistics for the fluorescence decays. Emission wavelengths from 494 to 530 nm were captured into a Single Molecule Detector Hybrid detector (HyD SMD). Fluorescence lifetimes were calculated using the Falcon (Fast Lifetime CONtrast) module (Leica LasX software) version 4.7.0. Selected areas of the images (+/- 3 cells/image) corresponding to either the PM or the TGN/EE were fitted by an n-exponential reconvolution fitting. The calculated IRF (Instrument response function) was included by setting the IRF background and shift to 0.

#### Protoplast swelling analysis

Root protoplasts were isolated from 7-day-old light-grown *Arabidopsis thaliana* seedlings grown vertically on nylon mesh over half-strength MS. Leaf protoplasts were isolated from 24-day-old light-grown plants cultivated in soil. Protoplast isolation was performed as previously described (Wendrich et al. 2020) with minor modifications. Briefly, roots and leaves were excised and incubated in protoplasting Solution B consisting of 1.5% (w/v) cellulase (Duchefa Biochemie C8003) and 0.1% (w/v) pectolyase (Duchefa Biochemie P8004) dissolved in Solution A (0.5 M mannitol, 10 mM CaCl₂, 20 mM MES, 20 mM KCl, pH 5.7). Samples were incubated for approximately 1 h at room temperature with gentle shaking (90 rpm). The released protoplasts were filtered through a 70-µm nylon cell strainer and centrifuged at 200 × g for 6 min. The protoplast pellet was gently resuspended in isotonic solution described hereafter.

Protoplast swelling experiments were performed as described previously with minor modifications (Moshelion et al. 2004; Shatil-Cohen et al. 2014). The isotonic and hypotonic solutions (10 mM KCl, 1 mM CaCl₂, 8 mM MES, pH 5.7) were adjusted with sorbitol to final osmolarities of 540 mM (isotonic) and 350 mM (hypotonic). Protoplast swelling was monitored by time-lapse imaging (1 image/s) on an inverted Leica DMIL microscope, with a 40X (NA 0.75) (1pixel = 0,209 µm) or a 20X (NA 0.3) (1pixel = 0,409 µm) dry objectives and equipped with a CCD video camera (Applied Imaging 4912-5010). Illumination was provided by transmitted light. Image acquisition was performed using the Scion Image software package (https://scion-image.software.informer.com/). Protoplasts were initially equilibrated in isotonic solution, and after 15 s the medium was switched to hypotonic solution to induce swelling. Note that while the hypotonic solution flow was switched on at 15 sec, it reached the bath only after a lag. Analysis of the protoplast surface area and volume changes were conducted using “Image explorer” and a “Protoplast Analyzer” plugins in the ImageJ software as described in (Shatil-Cohen et al. 2014). Curve morphologies for which the baseline values visually deviated strongly were discarded. Curves were smoothened using a walking average of 3 timepoints. Due to a variable swelling onset and occasional acquisition artifacts due to protoplast movement, analysis was performed after the stabilization phase, with early frames used only for baseline normalization.

The osmotic water permeability coefficient (P_f_) was determined using the Matlab *P_f_FIT* Program as explained in (Moshelion et al. 2004). Following P_f_ analysis, the quality of the output was visually screened to verify that the maximum slope was positioned in the linear area of the swelling phase. P_f_ values exceeding the mean ± 3 SD were considered as outliers and excluded.

### Chemical treatments

For the PI (Sigma, P4864) staining, 1 mg/ml of stock solution was diluted five times in water. Seedlings were dipped in the solution for 2 minutes and then rinsed in the water before imaging.

For the FM4-64 (Sigma, P4864) staining, 6-day-old seedlings were treated with 2 μM FM4-64 in 1/2 MS buffer for 15 minutes.

For the CHX (sigma, Lot #MKCL7692) and BFA (LC Laboratories, Lot #BBF-601) treatment, stock solutions of 50 mM in DMSO were made for both chemicals. Seedlings were pre-treated with 50 μM CHX for 30 minutes in 1/2 MS followed by co-treatment of CHX with 50 μM BFA for 1 hour in 1/2 MS.

For the BFA washout, the seedlings were treated with 50 μM BFA for 1 hour in 1/2 MS followed by rinsing in 1/2 MS for 2 hours.

### Image quantification

Quantifications of PM versus cytoplasm in the root meristem were performed using ImageJ and an in-house made script (Grones et al. 2022). Briefly, the script uses the PI staining in the red channel as a mask to detect and generate a PM and cytoplasm ROI region for the GFP channel. The top 5% mean grey value intensities in the green channel are measured in ImageJ. Two independent transgenic lines were imaged and quantified for each construct. For the quantification of rBiFC and the cell plate signal intensity following FM treatment, the PM or cell plate regions were selected by the active contour tool with a diameter of 8 pixels in ImageJ, and then the top 5% mean grey value intensities for YFP and RFP were measured in ImageJ. The quantifications of the Western blots were also performed in ImageJ. Bands of interest were selected by a square ROI, and their mean grey values were measured. Only images having fewer than 1% saturated pixels were used for quantification. The amount of saturated pixels was assessed using the histogram tool of ImageJ.

### Protein extraction and western blotting

#### Assessing PIP levels under normal growth conditions

7 day-old light-grown Arabidopsis seedlings were frozen in liquid nitrogen. Total proteins were extracted from around 100 mg of roots or leaves grinded in a ball mill and directly resuspended in 2x laemmli buffer and dithiothreitol (DTT) (2.5% SDS, 20% glycerol (v/v), 4% dithiothreitol, 0.01% bromophenol blue, 125 mM Tris HCl pH 6.8) (300 μl of buffer per 100 mg of tissue). Samples were heated at 65°C during 10 min, centrifuged 10 min at 20,000 × *g* at room temperature. Supernatants were collected and directly used for SDS-PAGE.

Equal protein amounts were loaded on a 4–20% SDS–PAGE TGX gel (BioRad, 4568093), and blotted on polyvinylidene difluoride (PVDF; BioRad). Membranes were blocked for 1 hour at room temperature using 5% skimmed milk. The blots were then incubated for 1 hour at room temperature with the endogenous antibody α-PIP1 (agrisera, AS09 487) or α-PIP2 (Santoni et al. 2003) (1:8000), which is followed by α-rabbit (Amersham ECL rabbit IgG, 1:15000). The anti-PIP1 antibody recognizes all PIP1 isoforms whereas anti-PIP2 recognizes PIP2;1, PIP2;2 and PIP2;3. Antibodies were applied in 2% skimmed milk and membranes were washed three times 10 minutes for each incubation. Detection of HRP chemiluminescence was performed using the SuperSignal^TM^ Chemiluminescent Substrate (Thermo Scientific) in a Chemidoc Touch Imaging system (Bio-Rad). Following image acquisition, the Image Lab software (Biorad) was used to quantify the signal obtained for a protein of interest. This signal was then normalized to the signal measured for the loading control.

#### Assessing PIP levels under drought-mimicking conditions

For sorbitol treatments, 7-day-old light-grown *Arabidopsis thaliana* seedlings were grown vertically on a nylon mesh. Seedlings were transferred to liquid ½ MS medium (mock) or ½ MS supplemented with 300 mM sorbitol for 90 min. After treatment, roots and leaves were carefully separated, and approximately 100 mg of tissue was collected for each sample. Total protein extraction and Western blotting were performed as described above for control conditions.

### Yeast two-hybrid and split-ubiquitin

5 μL aliquot Salm Sperm carrier DNA (Takara) and 40 μL yeast-competent cells were mixed for each construct, then 500 ng plasmid was added and incubated at 30°C for 15 minutes. 150μL PLI solution (for 15 ml PLI solution: 12 ml 50% PEG 3350 solution, 1.5 ml 1 M lithium acetate solution, and 1.5 ml sterile water) was added and mixed well by pipetting. The mixture was incubated at 30°C for 30 minutes followed by an incubation at 42°C for 15 minutes. Next, the samples were collected by centrifuging for 5 minutes at 3000g and the PLI was replaced with YPD. Samples were shaken for 2 hours at 200 rpm at 30°C. The transformants were plated on control plates (SD-L-T) and selective plates (SD-L-T-H, supplemented with 100 mM 3-Amino-1,2,4-triazole [3-AT]). The yeast minimal media and 3-AT are provided by Takara.

For split-ubiquitin assays, the haploid yeast strain THY.AP4 was co-transformed by electroporation with the Nub and Cub constructs of interest. Yeast colonies expressing the bait and prey constructs were recovered 72-96 hours after transfer to selective medium (CSM, -Leu−, Trp−) (Grefen et al. 2009). Protein expression was verified via immunoblotting using a rat monoclonal antibody against the hemagglutinin (HA) tag (Roche) and a rabbit polyclonal antibody against VP16 (Abcam), as previously described (Grefen et al. 2009). Protein loading on polyvinylidene fluoride (PVDF) membrane was revealed by Coomassie R250 staining (Goldman et al. 2016). Growth assays were performed as detailed in (Hachez et al. 2014). Yeast cultures were grown overnight on a selective medium. The next day, a dilution series (OD600 nm 0.5, 0.05, and 0.005) of the cultures was dropped onto interaction-selective (CSM, -Leu−, Trp−, Ade−, His−) medium containing 0, 100, 500 µM methionine to repress bait expression. Control plates CSM -Leu−, Trp− serve to verify that an equal yeast amount had been dropped. Yeast growth was recorded after incubation for 48 hours and 72 hours at 30°C.

### GFP pull-down

Pull-down was performed based on (Wendrich et al. 2017). Briefly, 7-day-old plate-grown Arabidopsis seedlings on 1/2 MS without sucrose were ground and harvested in liquid nitrogen, lysed in extraction buffer (50 mM Tris-HCl pH 7.5, 150 mM NaCl, 0.1% NP-40, 1× cOmplete Mini Protease Inhibitor Cocktail, Roche) and sonicated three times. After centrifugation two times for 15 minutes at 4°C at 18,000 rpm, 100 μl of anti-GFP μBeads (μMACS, Miltenyi) and supernatant were incubated for 2 hours at 4 °C. After that, the samples were run through μMACS Separator columns, which were equilibrated with extraction buffer. Columns were rinsed 4 times with extraction buffer, and then 2 times with ABC buffer (50 mM NH4HCO3 in H2O), and finally, samples were eluted with 95°C preheated ABC buffer.

Afterward, the samples were prepared for LC-MS/MS analysis. 1 μL 500 mM dithiothreitol (DTT) in ABC buffer was added to the beads and incubated for 2 hours at 60°C, which was followed by adding 1 μL 750 mM iodoacetamide (IAA) in ABC buffer to the samples. Then samples were incubated for 2 hours at room temperature in the dark. 1 μL 200 mM L-cysteine in the ABC buffer was added, and the next 1 μL sequence-grade trypsin was added before incubating for 16 hours at 20°C. The pH of the samples was adjusted to 3 by adding 10% TFA. All peptides in the samples were captured in the C18 resin. After the washing and elution steps, the samples were dried completely by vacuum drying.

Peptides were redissolved in 15 μl of loading solvent A (2% acetonitrile/0.1%TFA) of which 2 μl was injected on an Ultimate 3000 RSLCnano LC (Thermo Fisher Scientific) in-line connected to a Q Exactive mass spectrometer (Thermo Fisher Scientific) for LC-MS/MS analysis as previously reported(Persyn et al. 2024). The raw data were searched with MaxQuant (version 2.2.0.0) using the Andromeda search engine with the default settings, and a false discovery rate (FDR) set at 1% on peptide and protein level. To determine the significantly enriched proteins in the SCAMP5 bait sample compared to the *scamp5-1* control sample or in the SCAMP1 and SCAMP3 bait samples compared to the Col-0 control sample, Perseus (version 1.6.15.0) software was loaded with the proteingroups file from MaxQuant with reverse, contaminant, and only identified by site identifications removed. Also, the proteins with less than two valid values in at least one group were removed. LFQ intensities were log2 transformed and missing values were imputed from a normal distribution around the detection limit. A two-sided t-test was performed for pairwise comparison of the samples, and significantly enriched proteins were determined by permutation based FDR, using thresholds FDR = 0.05 and S0 = 1. The results are shown in the volcano plot in Figure 4 and the source data is available in Supplementary Table 2. The raw data and MaxQuant search data will be uploaded in the Pride database.

### AF modelling

SCAMP dimerization was modelled through AlphaFold2-Multimer (Evans et al. 2021) using a locally installed AlphaFold V2.3.2 version. The default parameters for MSA generation were used. 5 multimer predictions were run per model with a total of 5 models leading to a final number of 25 predictions per SCAMP dimer combination. Each model for the multimer prediction was calculated with a different random seed. The highest ranked model for each dimer combination was used for the analysis. Alphafold2-Multimer was run on a NVIDIA A5000 (24GB) GPU with CUDA version 12.1.

### Stomatal density measurements and dynamics

To measure stomatal density from soil-grown plants (21 °C with 70 ± 10% RH and 16 h light/8 h dark, 90 μmol m−2 s−1 photosynthetically active radiation from cool-white fluorescent tungsten tubes, Osram). assays were performed with modifications as described previously (Xu et al. 2025). The third leaf of 3-week-old Arabidopsis plants was cleared in cold 90% (v/v) acetone and mounted in 80% (v/v) lactic acid on a microscope slide. 4-6 non-overlapping photos from the abaxial epidermal cells at the centre of the leaf blade were taken using a Leica DMLB stereomicroscope, taking into account major veins. Stomatal density was determined by counting the number of stomata per unit area using the cell counter plugin of ImageJ. The average stomatal density of 4-6 images was calculated and displayed in Figure 6. Each genotype had 8 biological replicates.

Stomatal aperture assays were performed as previously described (Xu et al. 2025). Arabidopsis plants were grown and were manually equally watered similar to how plants were treated during our WIWAM experiment (Skirycz et al. 2011). Watering was subsequently halted. After 7 days of watering stop, the abaxial side of the third true leaf of three-week-old Col-0 and *scamp135^C^#1* plants was used for taking imprints. Moderate pressure was applied to combine the catalyst and polysiloxane (3M ESPE Express light body). The resulting mixture was spread onto the abaxial side of the leaf using toothpicks with minimal force, and once the mixture had solidified, the leaves were cut from the plants and the hardened imprint was removed. A layer of transparent nail varnish was applied to the hardened imprint and left to dry, creating a positive replica. The central region of these positive imprints was examined using a TM1000 scanning electron microscope (Hitachi). The acquired images were analysed and quantified using ImageJ. The length and width of the aperture of well-defined, properly imaged stomata was measured with Image J, and the ratio of aperture length to width was calculated.

## Data availability

The mass spectrometry data will be deposited to the ProteomeXchange Consortium (http://proteomecentral.proteomexchange.org) via the PRIDE partner repository (Perez-Riverol et al. 2019).

## Author contributions

Q.J., O.H. and D.V.D. designed the research and wrote the manuscript. All authors contributed to the final text. Q.J. generated most materials and performed SCAMP imaging analysis and mutant phenotyping analysis. M. VD. Performed RTPCR analysis and together with M. V. and Q.J. investigated the drought tolerance phenotype. A.R.F. performed split ubiquitin assays. O.H. performed western blotting, protoplast swelling experiments together with L.D. and APMS with the help of J.N. E.M. helped with confocal imaging. D.E. performed mass spectrometry data analysis. T.B. J. guided CRISPR/Cas9 technology. M.K and A.F.C. performed Alphafold modelling of SCAMPs. H.L. and I.D.S. performed and analysed stomatal measurements. D.K. and T.K.P. performed biochemistry experiments that in the end were omitted for simplicity. B.D.R., J.M.D., R.P., F.C. and D.V.D. were responsible for experimental design and research supervision.

## Acknowledgements

This work was supported by Funding from the Chinese Scholarship Council (CSC grant number 201906760018 to Q.J. and grant no. 202204910025 to H.L.), by the Special Research Fund of Gent University (BOF Covid prolongation funding to Q.J.) and by the Czech Science Foundation (grant number 22-35680M to R.P.). A.R.F. was supported by an individual Marie Sklodowska-Curie Action postdoctoral fellowship (Grant number: 101066810). Collaborative research in the Van Damme and Chaumont lab is supported by an FWO-WEAVE project (G0A7S24N to D.V.D. and to F.C.). The authors want to thank Mannon Waterland, Catherine Navarre and Adeline Courtoy (UCLouvain) for practical help, Cezary Waszczak (University of Helsinki) for constructive discussions, Marieke Dubois for help with the statistical quantification of the WIWAM data, the people of the VIB imaging core in Ghent for support with live cell imaging and the people from the VIB proteomics core for mass spectrometry measurements.

## Supplementary materials

**Supplementary Figure 1.**
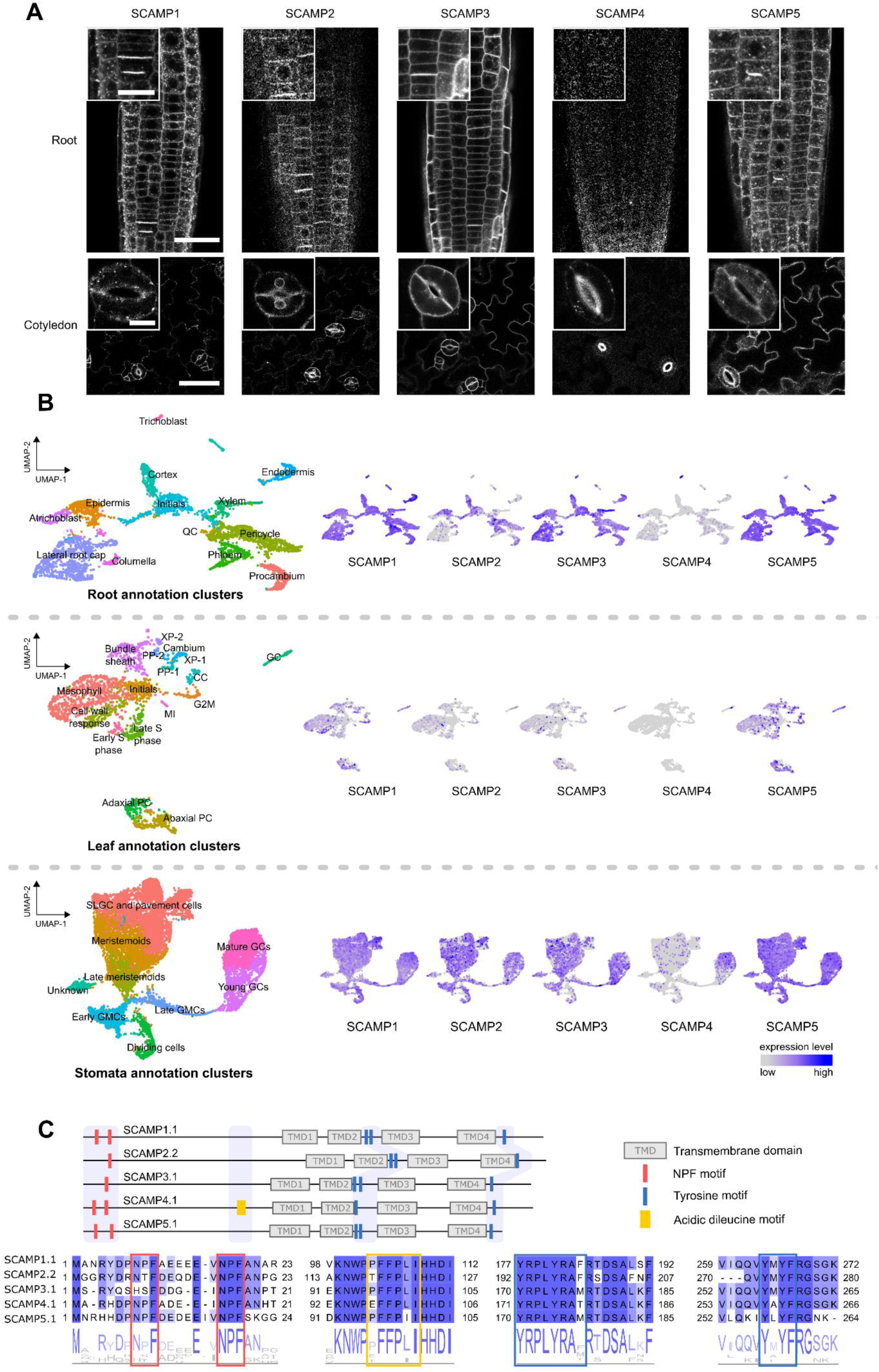
SCAMP family members have conserved trafficking motifs and are differentially expressed. **A)** Confocal images of roots and cotyledons of the five Arabidopsis SCAMPs tagged at the C-terminus with GFP and driven by their endogenous promotor sequence, with insets detail cell plate and stomata localization. Scale bar for the root pictures, 30 µm. Scale bar for the cell plate insets, 10 µm. Scale bar for the cotyledons, 50 µm. Scale bar for the stomata insets, 10 µm. **B)** Publicly available single-cell RNA sequencing results for the different SCAMPs in Arabidopsis roots, leaves, and stomatal lineage cells (Wendrich et al. 2020; Berrío et al. 2022; Kim et al. 2023). The color-coded UMAP plots on the left show the representative cell clusters for each tissue. QC: quiescent centre, XP-1: xylem parenchyma-1, XP-2: xylem parenchyma-2, PP-1: phloem parenchyma-1, PP-2: phloem parenchyma-2, CC: companion cell, GC: guard cell, MI: myrosin idioblast, Adaxial PC: adaxial pavement cell, Abaxial PC: abaxial pavement cell, SLGC: stomatal lineage ground cell, GMC: guard mother cell, GC: guard cell. The dot plots on the right depict the expression intensity of the different SCAMPs in cell sub clusters for each tissue according to the legend. **C)** Schematic representation of the transmembrane domains (TMDs), the NPF, tyrosine (Yxxϕ[I/L/M/F/V]), and the acidic dileucine ([D/E]xxxLL) motifs in Arabidopsis SCAMPs. The amino acid sequence alignments of the motifs are shown below. Numbers represent the amino acid positions of the motifs within the SCAMP family members. The TMD predictions were made by Phobius (Käll et al. 2004).

**Supplementary Figure 2.**
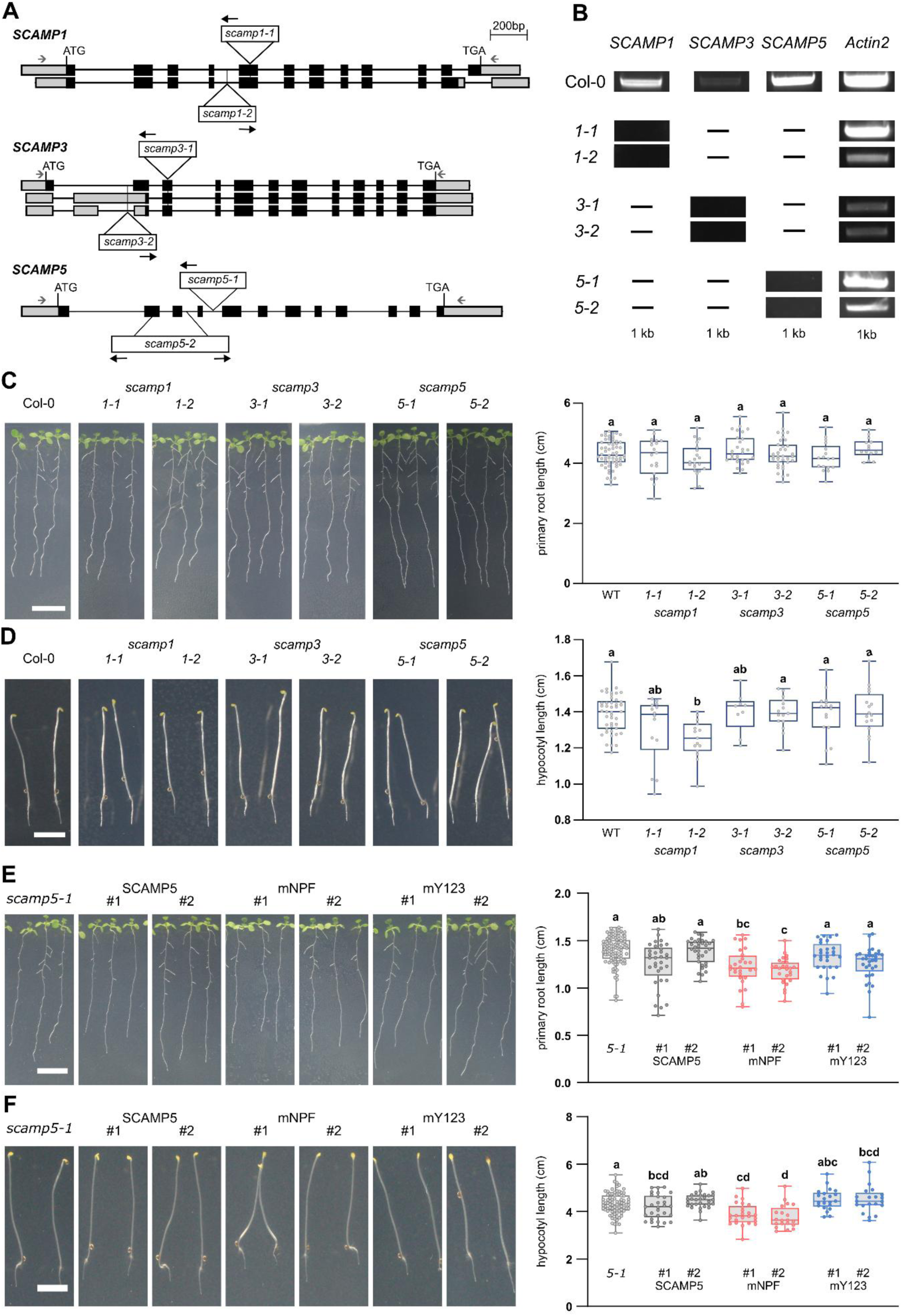
Characterization of the T-DNA mutants in *SCAMP1*, *SCAMP3* and *SCAMP5.* **A)** Analysed T-DNA insertion mutants of *SCAMP1*, *SCAMP3* and *SCAMP5* mapped on the different gene models. Exons are represented by black squares. Introns were indicated by black lines and untranslated regions are in grey squares. Insertion sites of T-DNA alleles identified in this study are indicated by white boxes. The black arrows indicate the orientations of insertions. Grey arrows represent primers for RT-PCR. **B)** RT-PCR analysis of Col-0 and single *scamp1, scamp3* and *scamp5* mutants. *ACTIN2* expression was used as control. The size of the amplicons is indicated. **C)** Representative images and box plot graphs depicting primary root length of 10-day-old Col-0 and single *scamp* mutants seedlings light-grown on 1/2 MS media without sucrose. Scale bar, 1 cm. 14 to 19 seedlings were quantified for each genotype. No significant difference in root length was found via a one-way ANOVA test (P>0,05). **D)** Representative hypocotyl images and box plot graphs depicting hypocotyl lengths of 4-day-old Col-0 and single *scamp* mutants seedlings dark-grown on 1/2 MS media without sucrose. Scale bar, 0,5 cm. 9 to 15 seedlings were quantified for each genotype. Different letters represent significant differences found via a one-way ANOVA test (P<0.05). **E)** Representative images and box plot graphs depicting primary root lengths of 10-day-old *scamp5-1* mutant seedlings as well as *scamp5-1* mutant seedlings expressing either SCAMP5 or motif-mutated forms of SCAMP5. Seedlings were light-grown on 1/2 MS media without sucrose. #1 and #2 represent independent lines. Scale bar, 1 cm. More than 20 seedlings were quantified for each genotype. Different letters represent significant differences following a one-way ANOVA test (P<0.05). **F)** Representative images and box plot graphs depicting hypocotyl lengths of 4-day-old *scamp5-1* mutant seedlings as well as *scamp5-1* mutant seedlings expressing either SCAMP5 or motif-mutated forms of SCAMP5. Seedlings were dark-grown on 1/2 MS media without sucrose. #1 and #2 represent two independent lines. Scale bar, 0,5 cm. More than 27 seedlings were quantified for each genotype. Different letters represent significant differences following a one-way ANOVA test (P<0.05). For each box plot, the bottom and top edges indicate the 25th and 75th percentiles. The central line is the median and whiskers extend to the full data range.

**Supplementary Figure 3.**
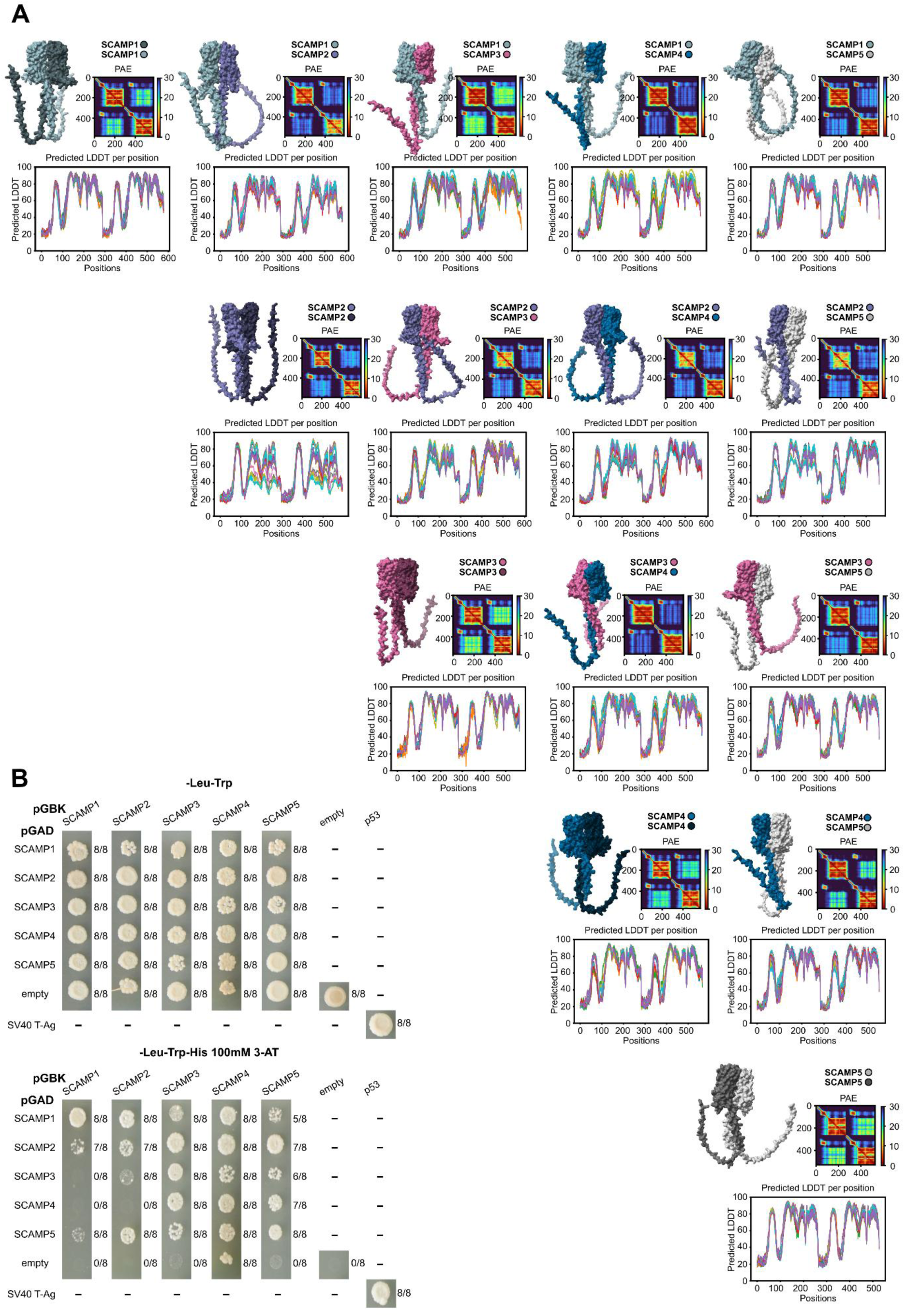
SCAMPs dimerize via their N-terminal domain. **A)** Overview of the best dimerization models from AlphaFold2 for all Arabidopsis SCAMP combinations. Prediction aligned error (PAE) plots that estimate the error of the position of each amino acid are shown for each combination. The predicted local distance difference test (LDDT) predicting local distance difference test for each combination covers 25 possible structures. **B)** Yeast two-hybrid assay combining the N-terminal cytoplasmic domains of all SCAMPs to check dimerization. Undiluted colonies expressing the designated constructs and empty vectors were spotted on -Leu-Trp plates and on -Leu-Trp-His plates with 100mM 3-Amino-1,2,4-triazole (3-AT). Growth was recorded after 72 h incubation. Numbers represent the amount of colonies out of a total of eight independent colonies for which growth was observed. The empty vector and The SV40 T-Ag and P53 combination, respectively, served as a negative and positive control. This experiment was repeated twice.

**Supplemental Figure 4:**
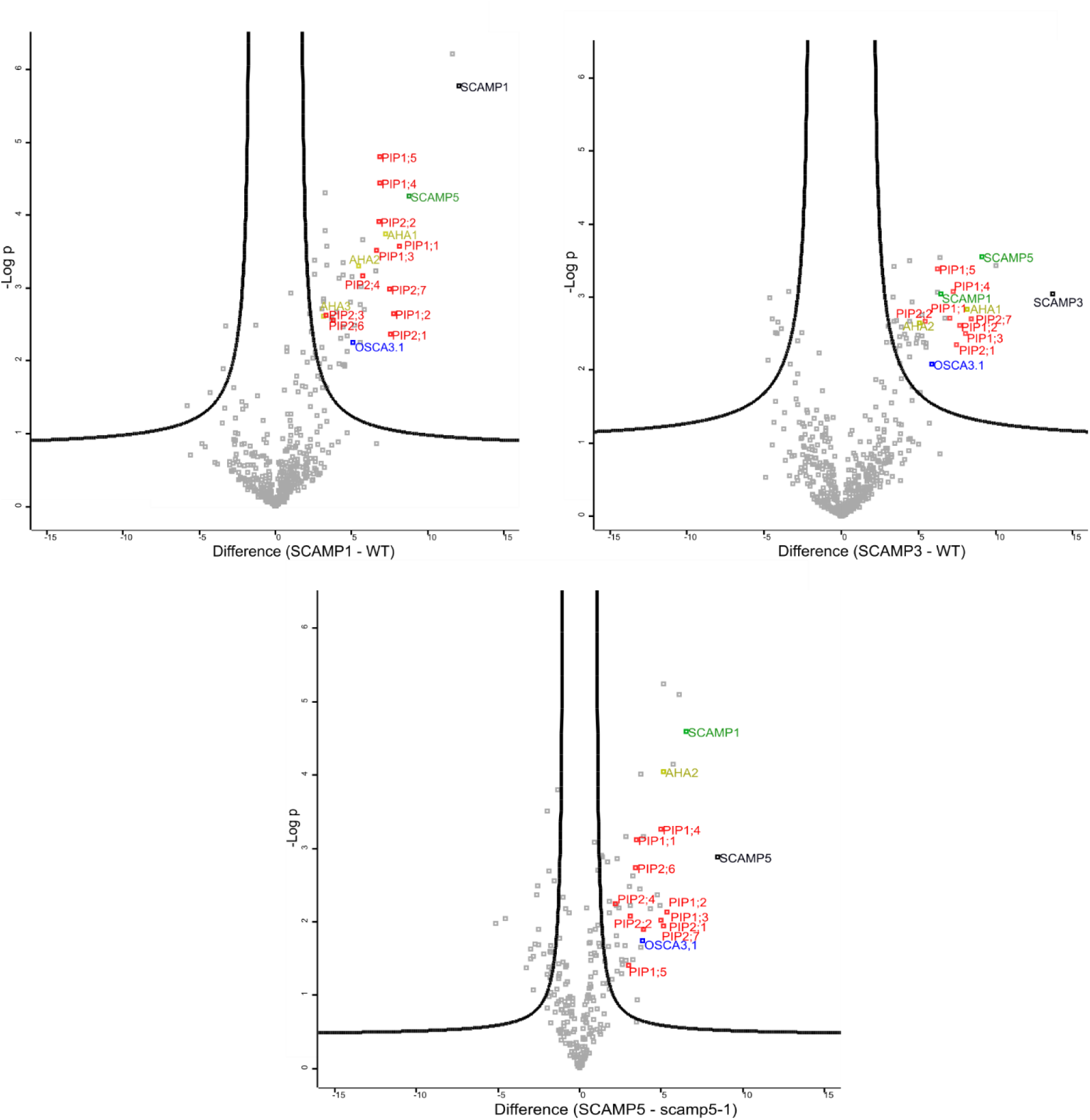
GFP-trap volcano plots of SCAMP1, SCAMP3 and SCAMP5. Volcano plots visualizing LFQ-based differential analysis of GFP-trap pulldown analyses from Arabidopsis seedlings. SCAMP1-GFP and SCAMP3-GFP expressed in the Col-0 background were used as bait proteins and compared with wild-type (Col-0) seedlings, whereas SCAMP5-GFP expressed in the *scamp5-1* (-/-) background was used as bait and compared with *scamp5-1* (-/-) mutant seedlings. Bait proteins are indicated in black. For pairwise comparison of the samples, a two-sided t-test was performed, and significantly enriched proteins were determined by permutation-based FDR, using FDR = 0.05 and S0 = 1 thresholds. Significantly enriched aquaporin family members (PIP), H+-ATPase (AHA), and the mechanically activated ion channel OSCA3.1 are highlighted in red, yellow and blue, respectively. Other SCAMP family members significantly enriched are indicated in green. This experiment was performed once with three technical repeats.

**Supplementary Figure 5.**
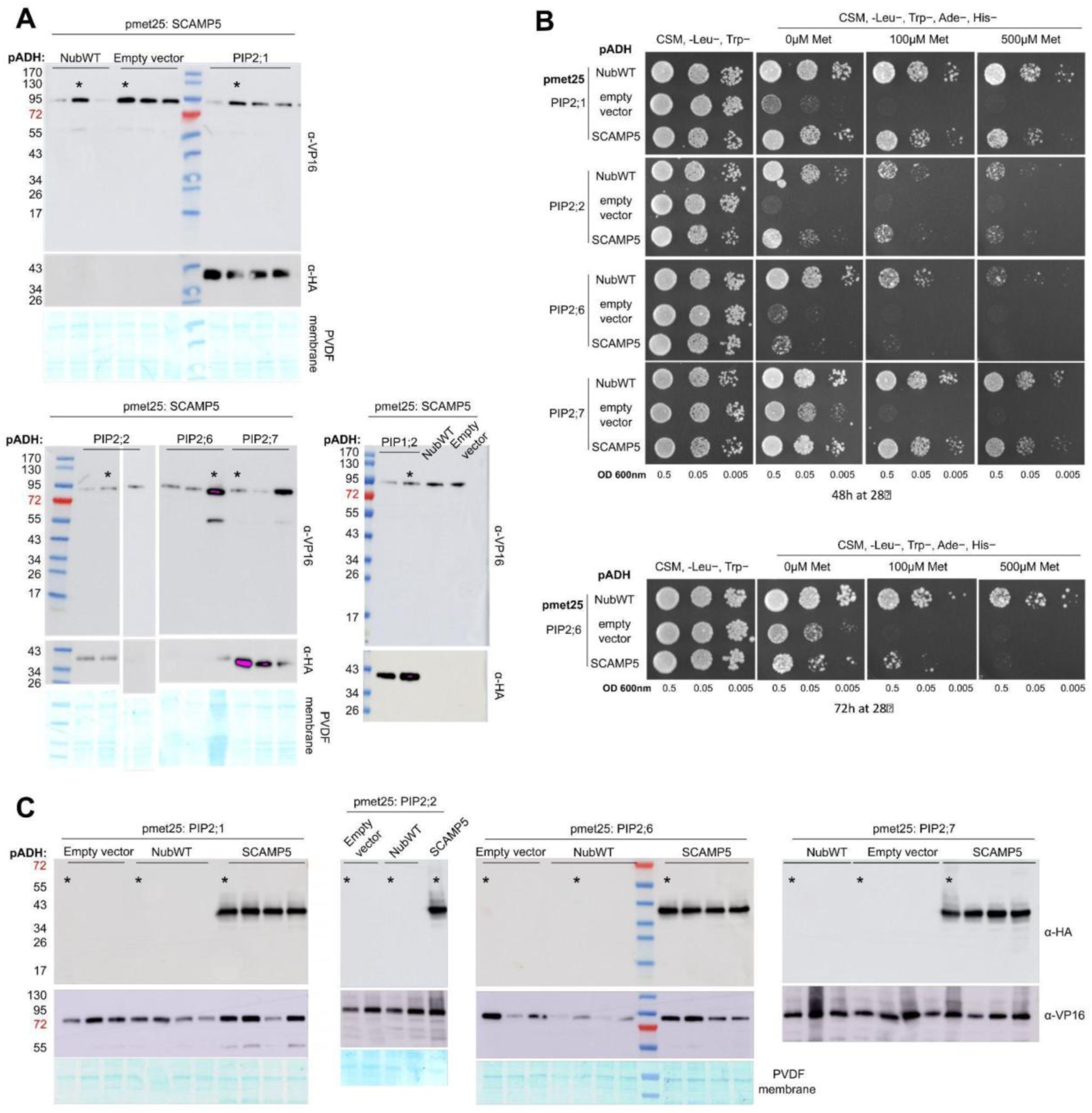
SCAMP5 interacts with PIP2 family members and yeast bait and prey expression analysis. **A)** Immunoblot analysis to verify bait and prey fusion protein expressions in transformed yeast prior to the split ubiquitin assays for Figure 4B. Only colonies expressing bait and preys were selected for the assays. Prey (PIP-NubA) expression was visualized using an anti-hemagglutinin (α-HA) antibody and the bait (SCAMP5-Cub-PLV) expression was revealed using an anti-VP16 antibody (α-VP16). NubWT does not contain a HA tag. The expected molecular weights of the proteins are: SCAMP5-Cub-PLV, 77 kDa; PIP-NubA, 40 kDa. Polyvinylidene difluoride (PVDF) membrane Coomassie R250 staining (bottom) was used as a loading control. **B)** Split ubiquitin assay to check if SCAMP5 can interact with various PIP2 isoforms when Cub and Nub tags are switched compared to Figure 4B. Yeast co-expressing the Met-repressible bait constructs PIP2;1/2/6/7-Cub-PLV and the prey constructs NubWT, Empty vector and NubG-SCAMP5 were spotted in a dilution series (OD 0.5, 0.05 and 0.005) onto synthetic medium (CSM, -Leu−, Trp−) and interaction-selective medium (CSM, -Leu−, Trp−, Ade−, His−) with increasing methionine concentrations to repress bait expression. Yeast growth was recorded after incubation for 48 h and 72 h. **C)** Immunoblot to verify bait and prey fusion protein expression in transformed yeast before the assay shown in panel B. The prey (NubG-SCAMP5) was revealed using an α-HA antibody and the bait (PIP2;1/2/6/7-Cub-PLV) expression was revealed using an α-VP16 antibody. NubWT does not contain HA-tag. The expected molecular weights of the proteins were: PIP2-Cub-PLV, 77 kDa; NubG-SCAMP5, 37.5 kDa. PVDF membrane Coomassie R250 staining (bottom) was used as loading control. The asterisks indicate those colonies that were used for the split-ubiquitin assays in this figure and in Figure 4B.

**Supplementary Figure 6.**
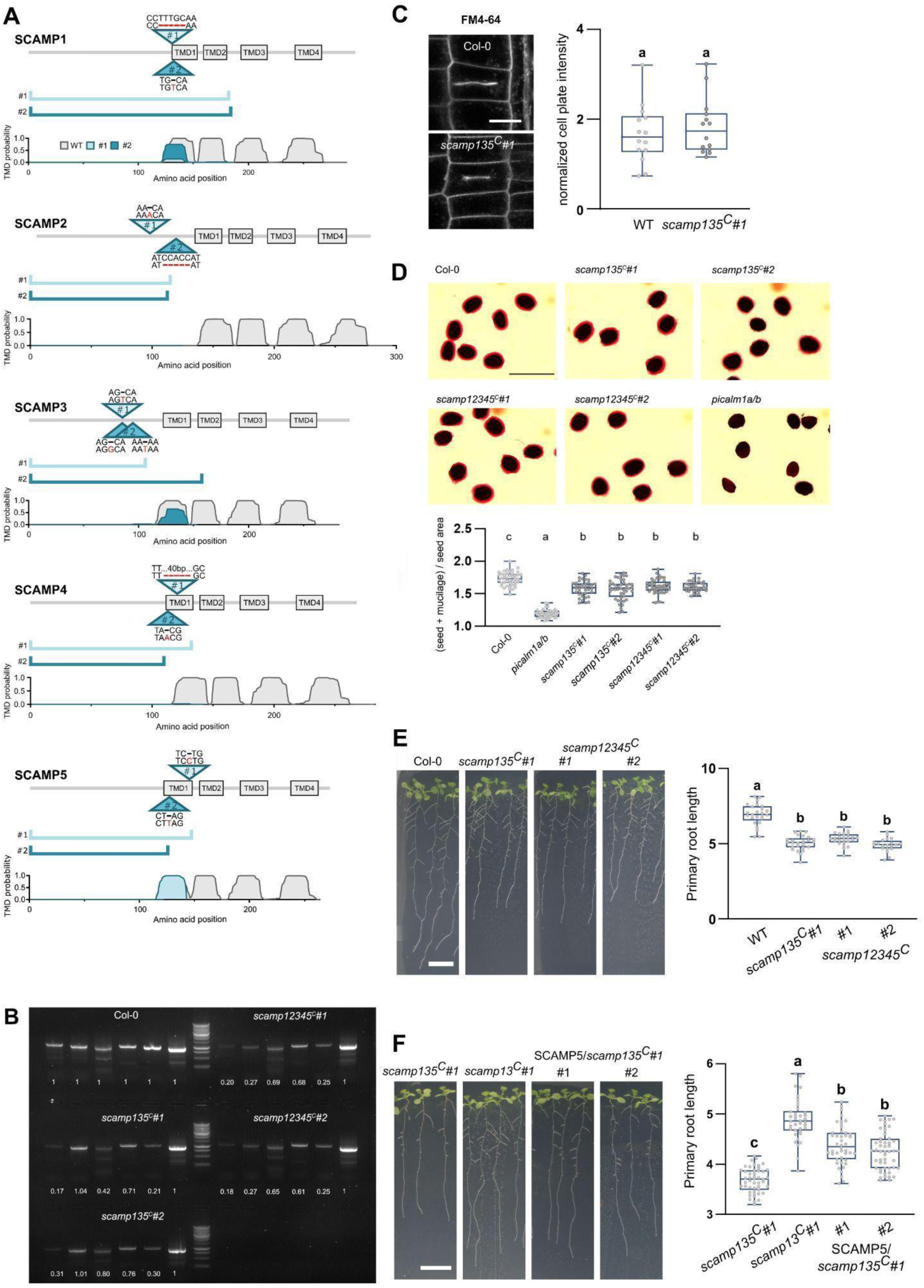
CRISPR-based *SCAMP* mutagenesis and characterization of the mutant lines. **A)** Schematic representation of CRISPR-induced mutations in all *SCAMPs.* The different triangles, indicate the gene editing positions that are light teal-coloured for #1 and dark teal-coloured for #2. The wild-type sequence and the mutated sequence are presented at the first and the second line respectively. Red hyphens represent deletions and red letters represent insertions. Transmembrane domains (TMDs) are shown as grey boxes. The teal lines depict theoretically translated SCAMPs after CRISPR-based gene editing, resulting in premature stop codons. The TMDs predictions of the wild-type SCAMPs (grey) are compared to the predictions for the CRISPR-edited alleles, light teal for #1, dark teal for #2 TMDs predictions were made by the Phobius software package (Käll et al. 2004). The triple *scamp135^C^#1* was created by combining the #1 edits in *SCAMP1*, *SCAMP3*, and *SCAMP5.* The triple *scamp135C#2* was created by combining the #2 edits in *SCAMP1*, *SCAMP3*, and *SCAMP5.* The quintuple *scamp12345^C^#1* and *scamp12345^C^#2* were generated by respectively combining the #1 edits or the #2 edits for *SCAMP2* and *SCAMP4* into the *scamp135C#1* triple mutant. **B)** Expression levels of the various *SCAMP* family members (*SCAMP1* to *SCAMP5* and *ACTIN2* as normalization control from left to right) determined by RT-PCR in WT (Col-0), two triple (scamp135^C^#1 and scamp135^C^#2) and two quintuple *scamp* mutant lines (*scamp12345^C^#1* and *scamp12345^C^#2*). The numbers represent average (n = 3) normalized intensity values of the amplified bands. Band intensities were normalized against the *ACTIN2* control. The numbers indicate the relative expression compared to WT levels. **C)** Representative images and box plot quantification of cell plate intensities of dividing root meristem cells of 6-day-old Col-0 and *scamp135^C^#1* seedlings stained with FM4-64. Seedlings were incubated in 1/2 MS with 2 µM FM4-64 for 15 min and then rinsed in 1/2 MS before imaging. Cell plate intensities were normalized to the intensities of the neighbouring plasma membrane. No significant difference was found between wild type and *scamp135^C^* triple mutant seedlings using a Student’s t-test (P>0,05). For each line, 14 cells from 7 seedlings were quantified. This experiment was performed once. **D)** Ruthenium red staining and quantification of the seed + mucilage versus seed area ratio of imbibed seeds from Col-0 as well as two independent *scamp135^C^* and *scamp12345^C^* lines. The *picalm1a/b* double mutant was used as a control. Scale bar, 1 mm. Different letters represent significant differences found by a one-way Anova including a Tukey multiple comparison test (p<0.0001). This experiment was performed twice. **E)** Representative images, and box plot quantification of primary root length comparing 13-day-old Col-0, *scamp135^C^#1* and *scamp12345^C^* (#1 and #2) seedlings light-grown on 1/2 MS media without sucrose. Scale bar, 1 cm. For each line, more than 20 seedlings were quantified. Different letters represent significant differences between wild-type, triple, and quintuple mutants as determined by a one-way ANOVA test (P<0.0001). This experiment was performed once. **F)** Representative images and box plot graph quantification of primary root length of 10-day-old *scamp135^C^#1*, *scamp13^C^#1* (combination of #1 edits in *SCAMP1* and *SCAMP3* from CRISPR editing), and two independent lines of *scamp135^C^#1* expressing SCAMP5-GFP. Seedlings light-grown on 1/2 MS media without sucrose. Scale bar, 1 cm. Different letters represent significant differences found by a one-way ANOVA test (P<0.05). This experiment was performed once. For each box in the box plot, the bottom and top edges indicate the 25th and 75th percentiles. The central line is the median and whiskers extend to the full data range.

**Supplementary Figure 7.**
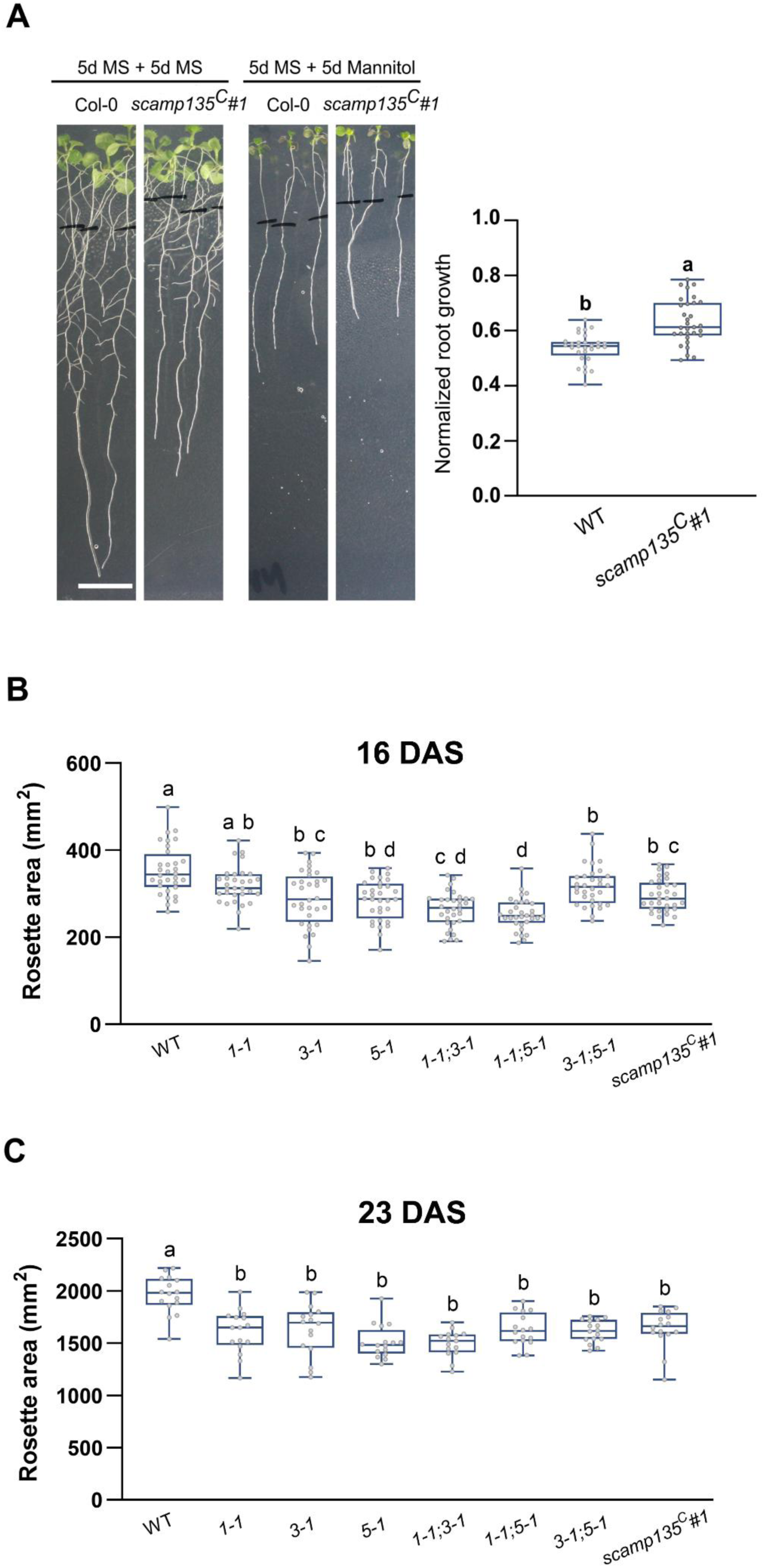
*scamp* mutants exhibit enhanced root growth under osmotic stress and reduced rosette area under well-watered conditions. **A)** Representative images and box plot quantification of root growth of WT and *scamp135^C^#1* mutants upon mannitol stress. 5-day-old seedlings were grown on 1/2 MS, then were transferred to 0 mM or 250 mM mannitol plates for another 5 days. The lines mark the position of the root after transfer. Scale bar, 1 cm. The graph depicts root growth on mannitol, normalized to the root growth under standard conditions. More than 20 seedlings were quantified for each condition. Different letters represent significant differences evaluated by a Student’s t-test (P<0.0001). This experiment was repeated twice. **B and C)** Box plots showing rosette area quantifications of WT and *scamp* mutants at 16 and 23 DAS under well-watered conditions. At 16 DAS (prior to drought treatment), rosette area was measured for 32 seedlings (B). A subset of 16 seedlings was maintained under well-watered conditions and re-assessed at 23 DAS (C). Different letters represent significant differences evaluated by a one-way ANOVA analysis including a Tukey’s multiple comparison test (0.0001<P<0.04) for (B) and (P>0.0001) for (C).

**Supplementary Table 1: used materials**

**Supplementary Table 2: MS results of the GFP-trap pulldowns from Arabidopsis seedlings, SCAMP5-GFP in *scamp5-1*(-/-) vs *scamp5-1*(-/-) seedlings and SCAMP1-GFP and SCAMP3-GFP in the Col-0 vs wild-type (Col-0) seedlings. Proteins were identified and quantified with MaxQuant and further analysed in Perseus. Significantly differing proteins were determined with a two-sided Student’s T-test and permutation-based FDR using thresholds FDR=0.05, S0=1.**

**Source file: summary of quantifications and source data.**

## References

Aoh QL, Castle AM, Hubbard CH, Katsumata O, and Castle JD. SCAMP3 negatively regulates epidermal growth factor receptor degradation and promotes receptor recycling. Mol Biol Cell. 2009:20(6):1816–1832. 10.1091/mbc.E08-08-0886

Arora D, Abel NB, Liu C, van Damme P, Yperman K, Eeckhout D, Vu LD, Wang J, Tornkvist A, Impens F, et al. Establishment of Proximity-Dependent Biotinylation Approaches in Different Plant Model Systems. Plant Cell. 2020:32(11):3388–3407. 10.1105/TPC.20.00235

Arora D and van Damme D. Motif-based endomembrane trafficking. Plant Physiol. 2021:186(1). 10.1093/plphys/kiab077

Asghari M, Abazari MF, Bokharaei H, Aleagha MN, Poortahmasebi V, Askari H, Torabinejad S, Ardalan A, Negaresh N, Ataei A, et al. Key genes and regulatory networks involved in the initiation, progression and invasion of colorectal cancer. Future Sci OA. 2018:4(3). 10.4155/fsoa-2017-0108

Berrío RT, Verstaen K, Vandamme N, Pevernagie J, Achon I, van Duyse J, van Isterdael G, Saeys Y, de Veylder L, Inzé D, et al. Single-cell transcriptomics sheds light on the identity and metabolism of developing leaf cells. Plant Physiol. 2022:188(2). 10.1093/plphys/kiab489

Bourdais G, McLachlan DH, Rickett LM, Zhou J, Siwoszek A, Häweker H, Hartley M, Kuhn H, Morris RJ, MacLean D, et al. The use of quantitative imaging to investigate regulators of membrane trafficking in Arabidopsis stomatal closure. Traffic. 2019:20(2):168–180. 10.1111/tra.12625

Brand SH, Laurie SM, Mixon MB, and David Castle J. Secretory Carrier Membrane Proteins 31-35 Define a Common Protein Composition among Secretory Carrier Membranes. Journal of Biological Chemistry. 1991:266(28):18949–18957.

Cai Y, Jia T, Lam SK, Ding Y, Gao C, San MWY, Pimpl P, and Jiang L. Multiple cytosolic and transmembrane determinants are required for the trafficking of SCAMP1 via an ER-golgi-TGN-PM pathway. Plant Journal. 2011:65(6). 10.1111/j.1365-313X.2010.04469.x

Castle A and Castle D. Ubiquitously expressed secretory carrier membrane proteins (SCAMPs) 1-4 mark different pathways and exhibit limited constitutive trafficking to and from the cell surface. J Cell Sci. 2005:118(16). 10.1242/jcs.02503

Chaumont F and Tyerman SD. Aquaporins: Highly Regulated Channels Controlling Plant Water Relations. Plant Physiol. 2014:164(4):1600–1618. 10.1104/PP.113.233791

Chen Q, Liu R, Wu Y, Wei S, Wang Q, Zheng Y, Xia R, Shang X, Yu F, Yang X, et al. ERAD-related E2 and E3 enzymes modulate the drought response by regulating the stability of PIP2 aquaporins. Plant Cell. 2021:33(8):2883–2898. 10.1093/plcell/koab141

Clough SJ and Bent AF. Floral dip: a simplified method for Agrobacterium -mediated transformation of Arabidopsis thaliana. The Plant Journal. 1998:16(6):735–743. 10.1046/J.1365-313X.1998.00343.X

Decaestecker W, Buono RA, Pfeiffer ML, Vangheluwe N, Jourquin J, Karimi M, van Isterdael G, Beeckman T, Nowack MK, and Jacobs TB. CRISPR-TSKO: A Technique for Efficient Mutagenesis in Specific Cell Types, Tissues, or Organs in Arabidopsis. Plant Cell. 2019:31(12):2868–2887. 10.1105/TPC.19.00454

Develtere W, Decaestecker W, Rombaut D, Anders C, Clicque E, Vuylsteke M, and Jacobs TB. Continual improvement of CRISPR-induced multiplex mutagenesis in Arabidopsis. Plant J. 2024:119(2):1158–1172. 10.1111/tpj.16785

Diering GH, Church J, and Numata M. Secretory Carrier Membrane Protein 2 Regulates Cell-surface Targeting of Brain-enriched Na+/H+ Exchanger NHE5. J Biol Chem. 2009:284(20):13892–13903. 10.1074/JBC.M807055200

Evans R, O’Neill M, Pritzel A, Antropova N, Senior A, Green T, Žídek A, Bates R, Blackwell S, Yim J, et al. Protein complex prediction with AlphaFold-Multimer. 2021. 10.1101/2021.10.04.463034

Fecht-Bartenbach J Von Der, Bogner M, Krebs M, Stierhof YD, Schumacher K, and Ludewig U. Function of the anion transporter AtCLC-d in the trans-Golgi network. Plant Journal. 2007:50(3). 10.1111/j.1365-313X.2007.03061.x

Fernández-Chacón R and Südhof TC. Novel SCAMPs Lacking NPF Repeats: Ubiquitous and Synaptic Vesicle-Specific Forms Implicate SCAMPs in Multiple Membrane-Trafficking Functions. Journal of Neuroscience. 2000:20(21):7941–50.

Fjorback AW, Müller HK, Haase J, Raarup MK, and Wiborg O. Modulation of the dopamine transporter by interaction with Secretory Carrier Membrane Protein 2. Biochem Biophys Res Commun. 2011:406(2):165–170. 10.1016/J.BBRC.2011.01.069

Fricke W and Knipfer T. Plant Aquaporins and Cell Elongation.. In. 10.1007/978-3-319-49395-4_5

Fujimoto M, Ebine K, Nishimura K, Tsutsumi N, and Ueda T. Longin R-SNARE is retrieved from the plasma membrane by ANTH domain-containing proteins in Arabidopsis. Proc Natl Acad Sci U S A. 2020:117(40). 10.1073/pnas.2011152117

Gadeyne A, Sánchez-Rodríguez C, Vanneste S, Di Rubbo S, Zauber H, Vanneste K, Van Leene J, De Winne N, Eeckhout D, Persiau G, et al. The TPLATE adaptor complex drives clathrin-mediated endocytosis in plants. Cell. 2014:156(4):691–704. 10.1016/J.CELL.2014.01.039

Garneau NL, Wilusz J, and Wilusz CJ. The highways and byways of mRNA decay. Nat Rev Mol Cell Biol. 2007:8(2). 10.1038/nrm2104

Goldman A, Harper S, and Speicher DW. Detection of proteins on blot membranes. Curr Protoc Protein Sci. 2016:2016. 10.1002/cpps.15

Grefen C and Blatt MR. A 2in1 cloning system enables ratiometric bimolecular fluorescence complementation (rBiFC). Biotechniques. 2012:53(5):311–314. 10.2144/000113941/ASSET/IMAGES/LARGE/FIGURE2.JPEG

Grefen C, Obrdlik P, and Harter K. The determination of protein-protein interactions by the mating-based split-ubiquitin system (mbSUS). Methods in Molecular Biology. 2009:479. 10.1007/978-1-59745-289-2_14

Grennan AK. Variations on a theme. Regulation of flowering time in arabidopsis. Plant Physiol. 2006:140(2). 10.1104/pp.104.900184

Grones P, De Meyer A, Pleskot R, Mylle E, Kraus M, Vandorpe M, Yperman K, Eeckhout D, Dragwidge JM, Jiang Q, et al. The endocytic TPLATE complex internalizes ubiquitinated plasma membrane cargo. Nat Plants. 2022:8(12):1467–1483. 10.1038/S41477-022-01280-1

Grubb LE, Derbyshire P, Dunning KE, Zipfel C, Menke FLH, and Monaghan J. Large-scale identification of ubiquitination sites on membrane-associated proteins in Arabidopsis thaliana seedlings. Plant Physiol. 2021:185(4):1483–1488. 10.1093/plphys/kiab023

Guo Z, Liu L, Cafiso D, and Castle D. Perturbation of a very late step of regulated exocytosis by a secretory carrier membrane protein (SCAMP2)-derived peptide. Journal of Biological Chemistry. 2002:277(38):35357–35363. 10.1074/jbc.M202259200

Hachez C, Laloux T, Reinhardt H, Cavez D, Degand H, Grefen C, De Rycke R, Inze D, Blatt MR, Russinova E, et al. Arabidopsis SNAREs SYP61 and SYP121 coordinate the trafficking of plasma membrane aquaporin PIP2;7 to modulate the cell membrane water permeability. Plant Cell. 2014:26(7):3132–3147. 10.1105/tpc.114.127159

Hachez C, Moshelion M, Zelazny E, Cavez D, and Chaumont F. Localization and quantification of plasma membrane aquaporin expression in maize primary root: A clue to understanding their role as cellular plumbers. Plant Mol Biol. 2006:62(1–2). 10.1007/s11103-006-9022-1

Hama Y, Kurikawa Y, Matsui T, Mizushima N, and Yamamoto H. TAX1BP1 recruits ATG9 vesicles through SCAMP3 binding. bioRxiv. 2023:2023.08.18.553817. 10.1101/2023.08.18.553817

Han C, Chen T, Yang M, Li N, Liu H, and Cao X. Human SCAMP5, a Novel Secretory Carrier Membrane Protein, Facilitates Calcium-Triggered Cytokine Secretion by Interaction with SNARE Machinery. The Journal of Immunology. 2009:182(5):2986–2996. 10.4049/jimmunol.0802002

He Z, Ma X, Zhu Q-H, Cheng S, Liu F, Zhang T, Zhang C, Li J, Xiong X, and Sun J. Genome-wide analysis of cotton SCAMP genes and functional characterization of GhSCAMP2 and GhSCAMP4 in salt tolerance. BMC Plant Biol. 2024:24(1):870. 10.1186/s12870-024-05571-x

Hubbard C, Singleton D, Rauch M, Jayasinghe S, Cafiso D, and Castle D. The Secretory Carrier Membrane Protein Family: Structure and Membrane Topology.

Hubert L, Cannata Serio M, Villoing-Gaudé L, Boddaert N, Kaminska A, Rio M, Lyonnet S, Munnich A, Poirier K, Simons M, et al. De novo SCAMP5 mutation causes a neurodevelopmental disorder with autistic features and seizures. J Med Genet. 2020:57(2). 10.1136/jmedgenet-2018-105927

Israel D, Lee SH, Robson TM, and Zwiazek JJ. Plasma membrane aquaporins of the PIP1 and PIP2 subfamilies facilitate hydrogen peroxide diffusion into plant roots. BMC Plant Biol. 2022:22(1):566. 10.1186/s12870-022-03962-6

Javot H, Lauvergeat V, Santoni V, Martin-Laurent F, Güçlü J, Vinh J, Heyes J, Franck KI, Schäffner AR, Bouchez D, et al. Role of a single aquaporin isoform in root water uptake. Plant Cell. 2003:15(2):509–22. 10.1105/tpc.008888

Javot H and Maurel C. The role of aquaporins in root water uptake. Ann Bot. 2002:90(3). 10.1093/aob/mcf199

Jiao X, Morleo M, Nigro V, Torella A, D’Arrigo S, Ciaccio C, Pantaleoni C, Gong P, Grand K, Sanchez-Lara PA, et al. Identification of an Identical de Novo SCAMP5 Missense Variant in Four Unrelated Patients With Seizures and Severe Neurodevelopmental Delay. Front Pharmacol. 2020:11. 10.3389/fphar.2020.599191

Käll L, Krogh A, and Sonnhammer ELL. A Combined Transmembrane Topology and Signal Peptide Prediction Method. J Mol Biol. 2004:338(5):1027–1036. 10.1016/J.JMB.2004.03.016

Karimi M, Depicker A, and Hilson P. Recombinational Cloning with Plant Gateway Vectors. Plant Physiol. 2007:145(4):1144–1154. 10.1104/pp.107.106989

Karimi M, De Meyer B, and Hilson P. Modular cloning in plant cells. Trends Plant Sci. 2005:10(3):103–105. 10.1016/j.tplants.2005.01.008

de Keijzer J, Kieft H, Ketelaar T, Goshima G, and Janson ME. Shortening of Microtubule Overlap Regions Defines Membrane Delivery Sites during Plant Cytokinesis. Curr Biol. 2017:27(4):514–520. 10.1016/j.cub.2016.12.043

Kelly BT, McCoy AJ, Späte K, Miller SE, Evans PR, Höning S, and Owen DJ. A structural explanation for the binding of endocytic dileucine motifs by the AP2 complex. Nature. 2008:456(7224). 10.1038/nature07422

Kim EJ, Zhang C, Guo B, Eekhout T, Houbaert A, Wendrich JR, Vandamme N, Tiwari M, Simon-Vezo C, Vanhoutte I, et al. Cell type–specific attenuation of brassinosteroid signaling precedes stomatal asymmetric cell division. Proc Natl Acad Sci U S A. 2023:120(36). 10.1073/pnas.2303758120

Kuromori T, Fujita M, Takahashi F, Yamaguchi-Shinozaki K, and Shinozaki K. Inter-tissue and inter-organ signaling in drought stress response and phenotyping of drought tolerance. Plant J. 2022:109(2):342–358. 10.1111/tpj.15619

Lampropoulos A, Sutikovic Z, Wenzl C, Maegele I, Lohmann JU, and Forner J. GreenGate - A novel, versatile, and efficient cloning system for plant transgenesis. PLoS One. 2013:8(12). 10.1371/journal.pone.0083043

Law AHY, Chow CM, and Jiang L. Secretory carrier membrane proteins. Protoplasma. 2012:249(2):269–283. 10.1007/s00709-011-0295-0

Lee HK, Cho SK, Son O, Xu Z, Hwang I, and Kim WT. Drought stress-induced Rma1H1, a RING membrane-anchor E3 ubiquitin ligase homolog, regulates aquaporin levels via ubiquitination in transgenic Arabidopsis plants. Plant Cell. 2009:21(2):622–41. 10.1105/tpc.108.061994

Lee U, Choi C, Ryu SH, Park D, Lee SE, Kim K, Kim Y, and Chang S. SCAMP5 plays a critical role in axonal trafficking and synaptic localization of NHE6 to adjust quantal size at glutamatergic synapses. Proc Natl Acad Sci U S A. 2021:118(2). 10.1073/pnas.2011371118

Liao H, Ellena J, Liu L, Szabo G, Cafiso D, and Castle D. Secretory carrier membrane protein SCAMP2 and phosphatidylinositol 4,5-bisphosphate interactions in the regulation of dense core vesicle exocytosis. Biochemistry. 2007:46(38):10909–10920. 10.1021/BI701121J

Liao H, Zhang J, Shestopal S, Szabo G, Castle A, and Castle D. Nonredundant function of secretory carrier membrane protein isoforms in dense core vesicle exocytosis. Am J Physiol Cell Physiol. 2008:294(3). 10.1152/ajpcell.00493.2007

Lin PJC, Williams WP, Luu Y, Molday RS, Orlowski J, and Numata M. Secretory carrier membrane proteins interact and regulate trafficking of the organellar (Na+,K+)/H+ exchanger NHE7. J Cell Sci. 2005:118(9):1885–1897. 10.1242/JCS.02315

Liu L, Guo Z, Tieu Q, Castle A, and Castle D. Role of secretory carrier membrane protein SCAMP2 in granule exocytosis. Mol Biol Cell. 2002:13(12). 10.1091/mbc.E02-03-0136

Liu M, Wang C, Ji Z, Lu J, Zhang L, Li C, Huang J, Yang G, Yan K, Zhang S, et al. Regulation of drought tolerance in Arabidopsis involves the PLATZ4-mediated transcriptional repression of plasma membrane aquaporin PIP2;8. Plant J. 2023:115(2):434–451. 10.1111/tpj.16235

Lykke-Andersen S, biology TJ-N reviews M cell, and 2015 undefined. Nonsense-mediated mRNA decay: an intricate machinery that shapes transcriptomes. Nat Rev Mol Cell Biol. 2015:16(11):665–677.

Meijering E, Jacob M, Sarria JCF, Steiner P, Hirling H, and Unser M. Design and Validation of a Tool for Neurite Tracing and Analysis in Fluorescence Microscopy Images. Cytometry Part A. 2004:58(2). 10.1002/cyto.a.20022

Melzer S, Lens F, Gennen J, Vanneste S, Rohde A, and Beeckman T. Flowering-time genes modulate meristem determinacy and growth form in Arabidopsis thaliana. Nat Genet. 2008:40(12). 10.1038/ng.253

De Meyer A, Grones P, and Van Damme D. How will I recognize you? Insights into endocytic cargo recognition in plants. Curr Opin Plant Biol. 2023:75. 10.1016/j.pbi.2023.102429

Moshelion M, Moran N, and Chaumont F. Dynamic changes in the osmotic water permeability of protoplast plasma membrane. Plant Physiol. 2004:135(4). 10.1104/pp.104.043000

Müller HK, Wiborg O, and Haase J. Subcellular redistribution of the serotonin transporter by secretory carrier membrane protein 2. J Biol Chem. 2006:281(39):28901–28909. 10.1074/JBC.M602848200

Mylle E, Codreanu MC, Boruc J, and Russinova E. Emission spectra profiling of fluorescent proteins in living plant cells. Plant Methods. 2013:9(1):1–8. 10.1186/1746-4811-9-10/FIGURES/3

Obudulu O, Mähler N, Skotare T, Bygdell J, Abreu IN, Ahnlund M, Gandla ML, Petterle A, Moritz T, Hvidsten TR, et al. A multi-omics approach reveals function of Secretory Carrier-Associated Membrane Proteins in wood formation of Populus trees. BMC Genomics. 2018:19(1). 10.1186/s12864-017-4411-1

Omasits U, Ahrens CH, Müller S, and Wollscheid B. Protter: Interactive protein feature visualization and integration with experimental proteomic data. Bioinformatics. 2014:30(6). 10.1093/bioinformatics/btt607

Park D, Lee U, Cho E, Zhao H, Kim JA, Lee BJ, Regan P, Ho WK, Cho K, and Chang S. Impairment of Release Site Clearance within the Active Zone by Reduced SCAMP5 Expression Causes Short-Term Depression of Synaptic Release. Cell Rep. 2018:22(12):3339–3350. 10.1016/j.celrep.2018.02.088

Perez-Riverol Y, Csordas A, Bai J, Bernal-Llinares M, Hewapathirana S, Kundu DJ, Inuganti A, Griss J, Mayer G, Eisenacher M, et al. The PRIDE database and related tools and resources in 2019: improving support for quantification data. Nucleic Acids Res. 2019:47(D1):D442–D450. 10.1093/nar/gky1106

Persyn F, Smagghe W, Eeckhout D, Mertens T, Smorscek T, De Winne N, Persiau G, Van De Slijke E, Crepin N, Gadeyne A, et al. A Nitrogen-specific Interactome Analysis Sheds Light on the Role of the SnRK1 and TOR Kinases in Plant Nitrogen Signaling. Mol Cell Proteomics. 2024:23(10):100842. 10.1016/j.mcpro.2024.100842

Pou A, Jeanguenin L, Milhiet T, Batoko H, Chaumont F, and Hachez C. Salinity-mediated transcriptional and post-translational regulation of the Arabidopsis aquaporin PIP2;7. Plant Mol Biol. 2016:92(6):731–744. 10.1007/s11103-016-0542-z

Renna L, Hanton SL, Stefano G, Bortolotti L, Misra V, and Brandizzi F. Identification and characterization of AtCASP, a plant transmembrane Golgi matrix protein. Plant Mol Biol. 2005:58(1). 10.1007/s11103-005-4618-4

Saeed B, Deligne F, Brillada C, Dünser K, Ditengou FA, Turek I, Allahham A, Grujic N, Dagdas Y, Ott T, et al. K63-linked ubiquitin chains are a global signal for endocytosis and contribute to selective autophagy in plants. Current Biology. 2023:33(7). 10.1016/j.cub.2023.02.024

Santoni V, Vinh J, Pflieger D, Sommerer N, and Maurel C. A proteomic study reveals novel insights into the diversity of aquaporin forms expressed in the plasma membrane of plant roots. Biochem J. 2003:373(Pt 1):289–96. 10.1042/BJ20030159

Shatil-Cohen A, Sibony H, Draye X, Chaumont F, Moran N, and Moshelion M. Measuring the osmotic water permeability coefficient (Pf) of spherical cells: isolated plant protoplasts as an example. J Vis Exp. 2014:(92):e51652. 10.3791/51652

Sheung KL, Cai Y, Hillmer S, Robinson DG, and Jiang L. SCAMPs highlight the developing cell plate during cytokinesis in tobacco BY-2 cells. Plant Physiol. 2008:147(4):1637–1645. 10.1104/pp.108.119925

Sheung KL, Ching LS, Hillmer S, Jang S, An G, Robinson DG, and Jiang L. Rice SCAMP1 defines clathrin-coated, trans-Golgi-located tubular-vesicular structures as an early endosome in tobacco BY-2 cells. Plant Cell. 2007:19(1):296–319. 10.1105/tpc.106.045708

Shimada T, Shimada T, Journal IH-TP, and 2010 undefined. A rapid and non-destructive screenable marker, FAST, for identifying transformed seeds of Arabidopsis thaliana. Plant Journal. 2010:61(3):519–528. 10.1111/j.1365-313X.2009.04060.x

Skirycz A, Vandenbroucke K, Clauw P, Maleux K, De Meyer B, Dhondt S, Pucci A, Gonzalez N, Hoeberichts F, Tognetti VB, et al. Survival and growth of Arabidopsis plants given limited water are not equal. Nat Biotechnol. 2011:29(3). 10.1038/nbt.1800

Smith S and De Smet I. Root system architecture: insights from Arabidopsis and cereal crops. Philos Trans R Soc Lond B Biol Sci. 2012:367(1595):1441–52. 10.1098/rstb.2011.0234

Sorieul M, Santoni V, Maurel C, and Luu DT. Mechanisms and Effects of Retention of Over-Expressed Aquaporin AtPIP2;1 in the Endoplasmic Reticulum. Traffic. 2011:12(4). 10.1111/j.1600-0854.2010.01154.x

Sparkes IA, Runions J, Kearns A, and Hawes C. Rapid, transient expression of fluorescent fusion proteins in tobacco plants and generation of stably transformed plants. Nat Protoc. 2006:1(4):2019–2025. 10.1038/nprot.2006.286

Toyooka K, Goto Y, Asatsuma S, Koizumi M, Mitsui T, and Matsuoka K. A mobile secretory vesicle cluster involved in mass transport from the golgi to the plant cell exterior. Plant Cell. 2009:21(4):1212–1229. 10.1105/tpc.108.058933

Ueda T, Uemura T, Sato MH, and Nakano A. Functional differentiation of endosomes in Arabidopsis cells. Plant Journal. 2004:40(5). 10.1111/j.1365-313X.2004.02249.x

Vadakekolathu J, Al-Juboori SIK, Johnson C, Schneider A, Buczek ME, Di Biase A, Pockley AG, Ball GR, Powe DG, and Regad T. MTSS1 and SCAMP1 cooperate to prevent invasion in breast cancer. Cell Death Dis. 2018:9(3). 10.1038/s41419-018-0364-9

Wang H, Tse YC, Law AHY, Sun SSM, Sun Y Bin, Xu ZF, Hillmer S, Robinsonand DG, and Jiang L. Vacuolar sorting receptors (VSRs) and secretory carrier membrane proteins (SCAMPs) are essential for pollen tube growth. Plant Journal. 2010:61(5):826–838. 10.1111/j.1365-313X.2009.04111.x

Wang X, Wu Y, Liu Z, Liu T, Zheng L, and Zhang G. The pip1s Quintuple Mutants Demonstrate the Essential Roles of PIP1s in the Plant Growth and Development of Arabidopsis. Int J Mol Sci. 2021:22(4). 10.3390/ijms22041669

Wang Y, Yang J, Jin H, Gu D, Wang Q, Liu Y, Zan K, Fan J, Wang R, Wei F, et al. Comparisons of physicochemical features and hepatoprotective potentials of unprocessed and processed polysaccharides from Polygonum multiflorum Thunb. Int J Biol Macromol. 2023:235. 10.1016/j.ijbiomac.2023.123901

Wendrich JR, Boeren S, Möller BK, Weijers D, and De Rybe L B. In vivo identification of plant protein complexes using IP-MS/MS.. In. Methods in Molecular Biology. 10.1007/978-1-4939-6469-7_14

Wendrich JR, Yang BJ, Vandamme N, Verstaen K, Smet W, Velde C Van de, Minne M, Wybouw B, Mor E, Arents HE, et al. Vascular transcription factors guide plant epidermal responses to limiting phosphate conditions. Science (1979). 2020:370(6518). 10.1126/science.aay4970

Wu TT and David Castle J. Tyrosine Phosphorylation of Selected Secretory Carrier Membrane Proteins, SCAMP1 and SCAMP3, and Association with the EGF Receptor.

Xu X, Liu H, Praat M, Pizzio GA, Jiang Z, Driever SM, Wang R, Van De Cotte B, Villers SLY, Gevaert K, et al. Stomatal opening under high temperatures is controlled by the OST1-regulated TOT3-AHA1 module. Nat Plants. 2025:11(1):105–117. 10.1038/s41477-024-01859-w

Yamada M, Matsuyama HJ, Takeda-Kamiya N, Sato M, and Toyooka K. Class II kinesin-12 facilitates cell plate formation by transporting cell plate materials in the phragmoplast. Nat Plants. 2025:11(2):340–358. 10.1038/s41477-025-01909-x

Yang B, Huang X, Zhang Y, Gao X, Ding S, Qi J, and Wang X. Genome-Wide Identification of Rubber Tree SCAMP Genes and Functional Characterization of HbSCAMP3. Plants (Basel). 2024:13(19). 10.3390/plants13192729

Yang Y, Qin M, Bao P, Xu W, and Xu J. Secretory carrier membrane protein 5 is an autophagy inhibitor that promotes the secretion of α-synuclein via exosome. PLoS One. 2017:12(7). 10.1371/journal.pone.0180892

Yavas I, Jamal MA, Din KU, Ali S, Hussain S, and Farooq M. Drought-Induced Changes in Leaf Morphology and Anatomy: Overview, Implications and Perspectives. Pol J Environ Stud. 2024:33(2). 10.15244/pjoes/174476

Yperman K, Papageorgiou AC, Merceron R, De Munck S, Bloch Y, Eeckhout D, Jiang Q, Tack P, Grigoryan R, Evangelidis T, et al. Distinct EH domains of the endocytic TPLATE complex confer lipid and protein binding. Nature Communications. 2021:12(1):1–11. 10.1038/s41467-021-23314-6

Zaarour N, Defontaine N, Demaretz S, Azroyan A, Cheval L, and Laghmani K. Secretory carrier membrane protein 2 regulates exocytic insertion of NKCC2 into the cell membrane. J Biol Chem. 2011:286(11):9489–9502. 10.1074/JBC.M110.166546

Zhang X, Sheng J, Zhang Y, Tian Y, Zhu J, Luo N, Xiao C, and Li R. Overexpression of SCAMP3 is an indicator of poor prognosis in hepatocellular carcinoma. Oncotarget. 2017:8(65). 10.18632/oncotarget.22665

Zhao H, Kim Y, Park J, Park D, Lee SE, Chang I, and Chang S. SCAMP5 Plays a critical role in synaptic vesicle endocytosis during high neuronal activity. Journal of Neuroscience. 2014:34(30):10085–10095. 10.1523/JNEUROSCI.2156-14.2014

